# Splicing modulators elicit global translational repression by condensate-prone proteins translated from introns

**DOI:** 10.1101/2020.11.23.393835

**Authors:** Jagat Krishna Chhipi Shrestha, Tilman Schneider-Poetsch, Takehiro Suzuki, Mari Mito, Khalid Khan, Naoshi Dohmae, Shintaro Iwasaki, Minoru Yoshida

**Affiliations:** Chemical Genomics Research Group, RIKEN Center for Sustainable Resource Science, Wako, Saitama 351-0198, Japan; Department of Biotechnology, Graduate School of Agricultural and Life Sciences, The University of Tokyo, Bunkyo-ku, Tokyo 113-8657, Japan; Biomolecular Characterization Unit, Technology Platform Division, RIKEN Center for Sustainable Resource Science, Wako, Saitama 351-0198, Japan; RNA Systems Biochemistry Laboratory, RIKEN Cluster for Pioneering Research, Wako, Saitama 351-0198, Japan; Department of Computational Biology and Medical Sciences, Graduate School of Frontier Sciences, The University of Tokyo, Kashiwa, Chiba 277-8561, Japan; AMED-CREST, Japan Agency for Medical Research and Development, Wako, Saitama 351-0198, Japan; Collaborative Research Institute for Innovative Microbiology, The University of Tokyo, Bunkyo-ku, Tokyo 113-8657, Japan

## Abstract

Chemical splicing modulators that bind to the spliceosome have provided an attractive venue for cancer treatment. Splicing modulators induce accumulation and subsequent translation of a subset of intron-retained mRNAs. Yet, the biological effect of proteins containing translated intron sequences remains unclear. Here we identified a number of truncated proteins generated upon treatment with the splicing modulator spliceostatin A (SSA) using genome-wide ribosome profiling and bio-orthogonal non-canonical amino-acid tagging (BONCAT) mass spectrometry. A subset of these truncated proteins has intrinsically disordered regions, forms insoluble cellular condensates, and triggers the proteotoxic stress response through JNK phosphorylation, thereby inhibiting the mTORC1 pathway. In turn, this reduces global translation. These findings indicate that creating an overburden of condensate-prone proteins derived from introns represses translation and prevents further production of harmful truncated proteins. This mechanism appears to contribute to the antiproliferative and proapoptotic activity of splicing modulators.

## Introduction

Splicing, the removal of intervening sequences from pre-mRNAs, is an essential step in mRNA maturation in eukaryotes to maintain correct gene expression. Inadequate splicing is associated with a variety of diseases and tumorigenesis ^1–3^. Clinical genome sequencing revealed that mutations in spliceosome genes such as *SF3B1*, *SRSF2*, and *U2AF1* occur at surprisingly high frequencies in hematological malignancies, including myelodysplastic syndromes (MDS) and chronic lymphocytic leukemia (CLL) ^4, 5^. Broadly, aberrant splicing patterns are frequently seen in cancers ^6^. Even splicing pattern switching of a single gene (pyruvate kinase PKM) may lead to tumorigenesis ^7^.

Due to the tight associations between tumors and splicing dysregulation, recently identified chemical modulators of splicing have drawn considerable interest in their therapeutic potential ^8^. Since the discovery of natural products FR901464 and pladienolide B (PlaB) as small molecules that specifically bind and inhibit SF3B1 — a component of the SF3B subcomplex of the U2 snRNP ^9–11^, a variety of structurally related splicing modulators, such as spliceostatin A (SSA), sudemycin, meayamycin, and E7107, have been developed. As candidates for cancer therapeutics (see review by ^12, 13^), these splicing modulators show the capability to suppress cancer cells expressing mutant spliceosomal proteins in both *in vitro* and in animal models ^5, 14–16^. In particular, H3B-8800, an orally available molecule ^16^, has begun clinical trials for treating both solid tumors and leukemias bearing spliceosome mutations. Even in the absence of splicing factor mutations, chemical splicing modulators appear to specifically induce apoptosis in a wide variety of tumor cells ^17–20^.

A number of mechanisms by which splicing modulators inhibit tumor cell survival and proliferation have been proposed: (1) synthetic lethality with splicing factor mutations ^16, 21, 22^, (2) excess demand of spliceosome activity in MYC-activated tumor cells ^23^, (3) splicing perturbation of BCL2 family antiapoptotic genes ^24–26^, and (4) inhibition of angiogenesis by downregulating vascular endothelial growth factor (VEGF) expression in malignant tumors ^27–29^. However, these explanations only apply to certain types of cancer. The generalized rationale for how splicing modulators suppress tumor growth remains unclear.

The outcome of splicing modulator treatment is not only limited to the downregulation of spliceosome activity but alters downstream mRNA processing. Inhibition of splicing normally induces drop-off and/or elongation arrest of RNA polymerase II (Pol II) via dephosphorylation of Ser2 in the C-terminal domain (CTD) ^30, 31^. Moreover, mRNAs containing introns are retained inside the nucleus ^9, 32–34^, followed by degradation via the 3’-5’ riboexonuclease exosome ^35^. However, a subset of mRNAs still leak into the cytoplasm ^9, 33, 34^. This fraction of mRNAs is usually degraded via the nonsense-mediated decay (NMD) pathway, which recognizes premature termination codons (PTCs) inside intronic sequences (reviewed in ^36^).

Despite all these quality control mechanisms, a substantial fraction of intron-retained mRNAs escapes surveillance and thereby yields truncated, incomplete proteins with extensions from introns ^9, 37, 38^. We previously observed that FR901464 and SSA induce the production of truncated forms (we denote the truncated protein as * following earlier nomenclature ^9^) of the tumor suppressor protein p27 (p27*) and the NF-κB signaling inhibitory protein IκBα (IκBα*)^9^. This suggests the possibility that a significant number of functionally active incomplete proteins are generated upon splicing modulator treatment. Although short transcripts with weaker 5′ splice sites are prone to leak from the nucleus ^33^, the generation of truncated proteins from such leaked mRNA by splicing modulators has not yet been systematically explored.

To address this issue, we implemented a global approach combining transcriptome, translatome, and proteome analyses upon chemical splicing perturbation. We found that under SSA treatment, a wide array of transcripts experience intron translation leading to proteins possessing intrinsically disordered and condensate-prone regions. This condensation activates the proteotoxic stress response via JNK phosphorylation and in turn inhibits the mTORC1 pathway. Inhibition of mTORC1-mediated translation activation significantly reduces the output of protein biosynthesis. Our results present an unexpected property of deleterious proteins originating from erroneous mRNA processing.

## Results

### SSA induces widespread intron retention and intron translation

To globally survey the truncated proteins generated by chemical splicing modulation beyond p27* and IκBα*, we performed simultaneous ribosome profiling and mRNA sequencing in the presence and absence of SSA (Extended Data Fig. 1A). Ribosome profiling is a powerful method based on deep sequencing of ribosome-protected mRNA fragments and provides the best overview of translation dynamics at a subcodon resolution ^39, 40^, enabling the global identification of translated introns. Our data displayed high experimental reproducibility (Extended Data Fig. 1B) and sample quality, including the expected size of ribosome footprints (Extended Data Fig. 1C) and three-nucleotide periodicity along the coding sequence (CDS) (Extended Data Fig. 1D). Simultaneously, we performed sequencing of cellular RNAs (RNA-Seq) to evaluate the occurrence of splicing changes under SSA treatment.

**Fig. 1:**
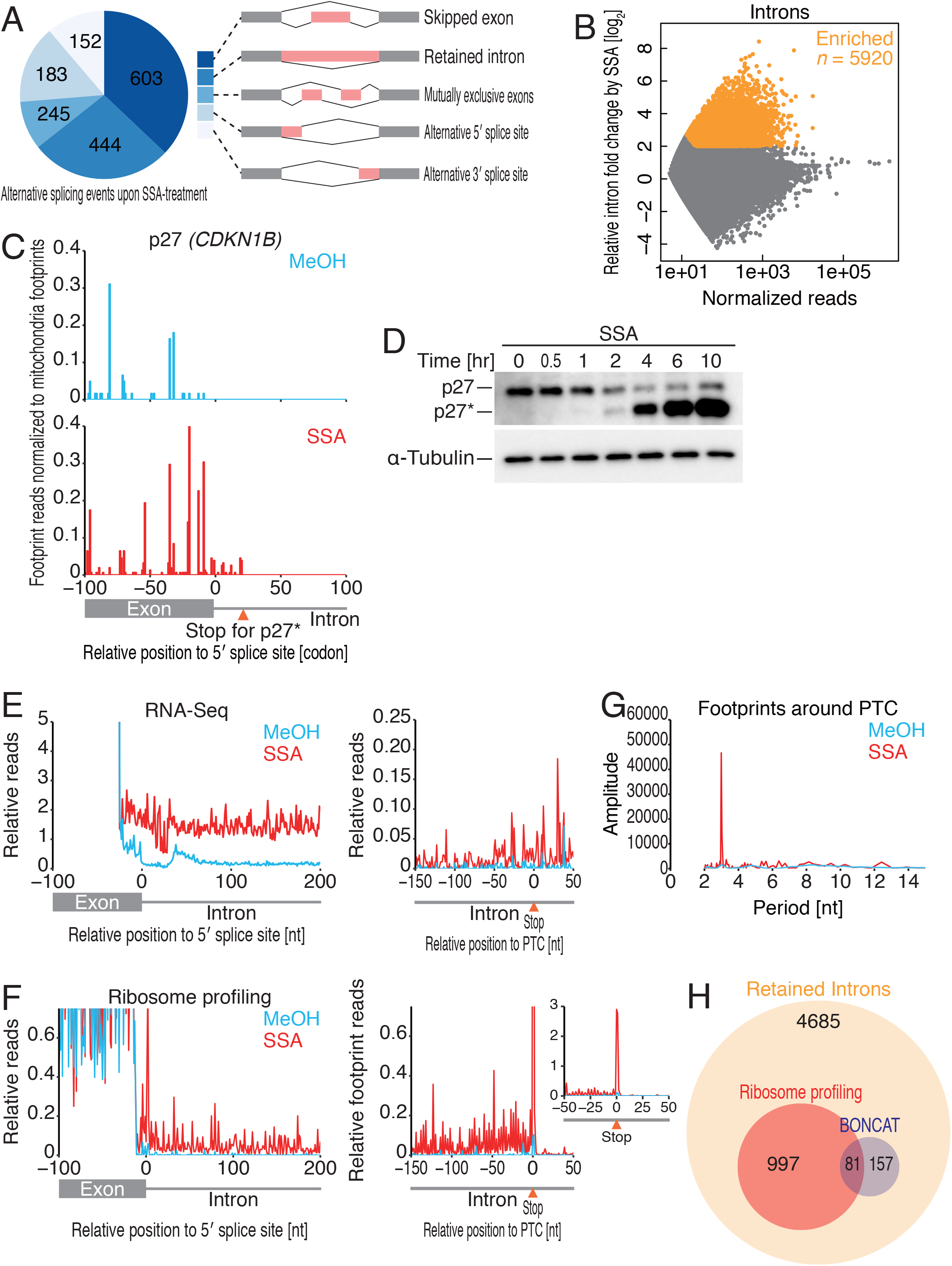
Retained introns that emerged upon splicing modulation are extensively subjected to translation. (A) SSA-induced transcriptome-wide splicing alterations, analyzed using the MISO framework ^87^. Exon skipping and intron retention were the most common effects observed in the presence of SSA. (B) MA (log ratio vs. mean average) plot for intron enrichment in mRNAs affected by SSA, displayed as relative intron fold change. Significantly enriched introns [false discovery rate (FDR) < 0.01, log_2_-fold change ≥ 2] are highlighted in orange. (C) Ribosome footprint accumulation in the intron of p27 (*CDKN1B)* under splicing inhibition. Reads were normalized to the sum of mitochondrial footprints. (D) Western blot for p27 and p27* in the HeLa S3 cell lines treated with MeOH solvent or 100 ng/ml SSA for different time periods. (E and F) Meta-gene analysis of translated introns in RNA-Seq reads (E) and ribosome footprints (F) relative to the 5’ splice site (left) and the PTC (right). The reads were normalized to the sum of exonic RNA reads from 100 nucleotides upstream from the 5’ splice site. Fig. 1F contains the zoomed out inset to account for the height of the peak at the PTC. (G) Discrete Fourier transform of ribosome footprint reads to visualize the periodicity around the PTC. (H) Venn diagram showing the total number of retained introns and the fractions of translated introns detected by ribosome profiling and BONCAT. The reference database was prepared from the *in silico* translation of retained introns. For BONCAT, peptides spanning the exon-intron junction were considered.

Among the different alternative splicing events observed in the RNA-Seq data, SSA induces exon skipping and intron retention (Fig. 1A), confirming the results of previous studies on SF3B inhibitors ^9, 33, 41, 42^. Analysis of global intronic reads (see Materials and Methods for details) ensured that a number of transcripts contained retained introns (Fig. 1B). We define this subgroup of introns as “retained introns” and their source mRNAs as “intron-retained mRNAs”. We confirmed the findings by RT-PCR of a representative transcript (*DNAJB1*) that clearly showed intron retention upon SSA treatment (Extended Data Fig. 1E).

Ribosome profiling demonstrated that a large number of the retained introns did reach the translation machinery. Intronic ribosome footprints on the p27 mRNA were found until the first in-frame stop codon (Fig. 1C), corresponding to the truncated protein seen by Western blotting (Fig. 1D). Similarly, intron translation was widely observed across the transcriptome. Meta-gene analysis centered on the exon-intron junction showed a substantial increase in intronic reads upon SSA treatment, both in mRNA reads and in ribosome footprints (Fig. 1E and 1F, left). Of the 5920 retained introns detected by RNA-Seq in the presence of SSA, we observed active translation of 1078 intronic sequences (Extended Data Table 1). Moreover, we found that the number of footprints dropped off at the first in-frame stop codon (Fig. 1F, right). Ribosome footprints upstream of the PTC showed 3-nt periodicity when analyzed by discrete Fourier transform, which indicates active translation from the retained introns (Fig. 1G).

In line with previous reports, we observed that a significant fraction of mRNAs containing intronic sequences escaped NMD surveillance ^9, 37, 38^. The intron-translated mRNAs should have been targeted by the NMD pathway, since the distance from the intronic PTC to the downstream exon-exon junction exceeded ∼50-55 nucleotides (nt) the minimal distance that triggers NMD (Extended Data Fig. 1G, right) ^43^.

We further analyzed the products of intron translation by a proteome approach. Since the pre-existing proteins perturb the detection of the translated peptides from intron-retained mRNAs that are evoked in the time frame of SSA treatment (6 h), here we set out to enrich the newly synthesized protein during the SSA treatment by bio-orthogonal non-canonical amino-acid tagging (BONCAT). This technique is based on metabolic labeling of the newly synthesized proteins by non-canonical amino acid homopropargylglycine (HPG) that allows cycloaddition of azide-biotin by “click chemistry”, enrichment of the biotinylated proteins by streptavidin beads, and subsequent detection of *de novo* synthesized proteins by mass spectrometry ^44, 45^. Creating a database of predicted chimeric introns in frame with the translation start from the 5920 retained introns (Fig. 1B), BONCAT detected a substantial number of stable chimeric proteins (n = 238) (Extended Data Table 2) in the lysate prepared by SSA treatment (Fig. 1H). The number of observed chimeric intron proteins is comparatively smaller (∼20% only) than the number observed in ribosome profiling, likely due to the predominant presence of peptides from full-length proteins (2852 proteins detected) even in the BONCAT strategy and global repression of protein synthesis (see below).

Taken together, our results demonstrate that SSA treatment leads to the production of diverse chimeric proteins containing both exon- and intron-derived sequences.

### Characteristics of intron-translated transcripts and proteins

In the course of characterizing the intron-translated transcripts, we observed some general properties applying to the majority of predicted chimeric intron-translated peptides and their source transcripts. A majority of the translated introns (71%) were derived from the first intron (Extended Data Fig. 2A, right), implying that translation halts on the PTC (Fig. 1F) and does not progress downstream (Extended Data Fig. 2A, left). These chimeric intron-translated proteins have a median length of 94 amino acids (Fig. 2A).

**Fig. 2:**
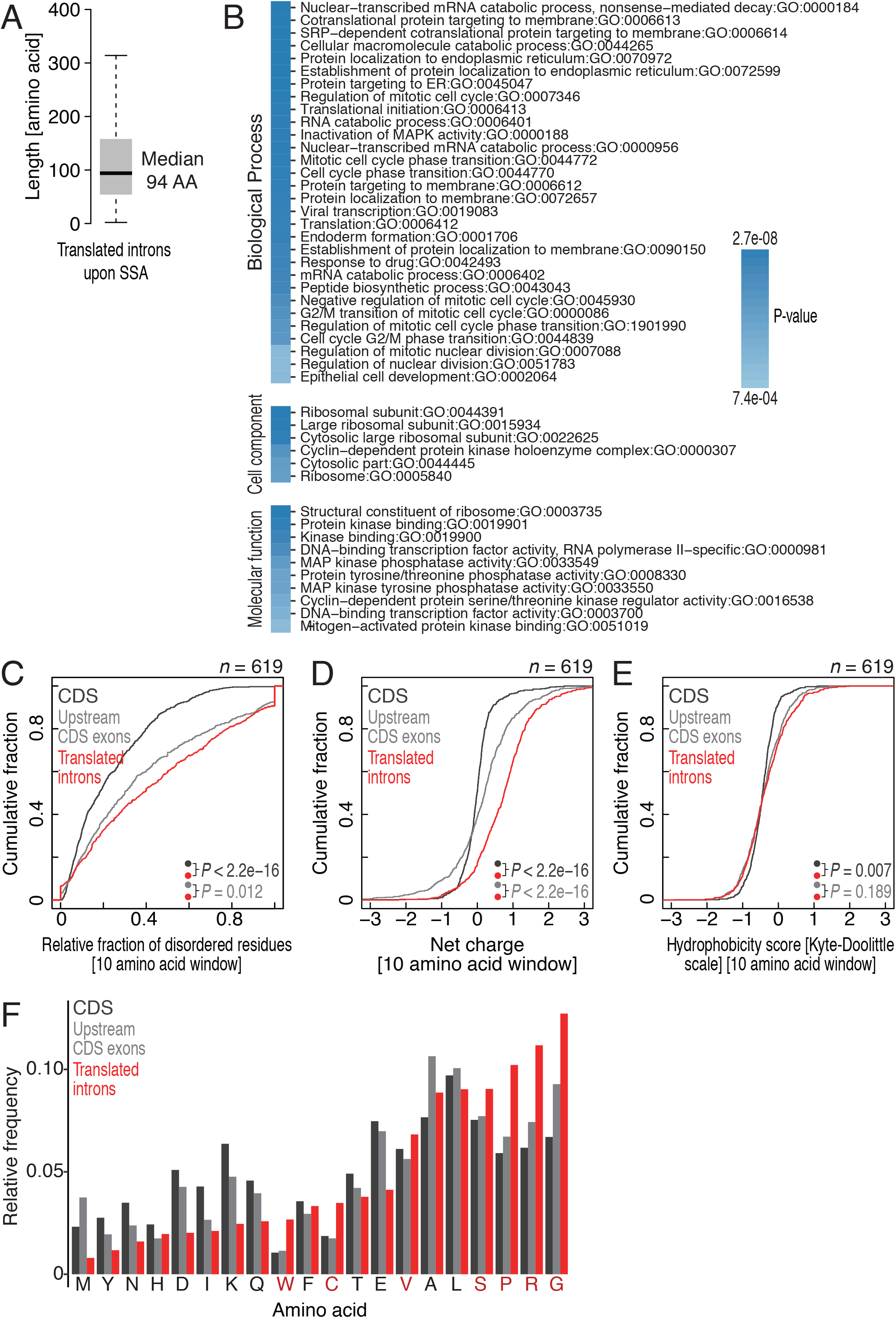
Characterization of translated introns. (A) Predicted length of translated introns upon SSA. The median length is shown. (B) Gene ontology (GO) terms of source genes of translated introns. Color indicates the statistical value. (C-E) Cumulative distribution of relative disorderness (IUPred2A score) (C), net charge (Lehninger pKa scale) (D), and hydrophobicity score (Kyte-Doolittle scale) (E) of amino acid residues in translated introns, upstream CDS exons, and full CDS. Ten amino acid windows were considered. (F) Relative frequency of amino acids present in either translated introns, upstream CDS exons, or CDS. Amino acids overrepresented in translated introns are highlighted in red. In C-E, the significance was calculated by Wilcoxon’s test.

To determine whether these intron-translated mRNAs have particular cellular functions, we conducted gene ontology analysis. Genes pertaining to mRNA catabolism, protein targeting to the membrane, and the cell cycle were significantly enriched (Fig. 2B).

In contrast to full-length proteins, truncated peptides may lack well-defined and stable secondary and tertiary structures. IUPred2A ^46^ prediction of intrinsically disordered regions (IDRs) indicated that translated introns have significantly higher levels of IDRs than upstream CDS exons or the full-length CDS of the same transcript (Fig. 2C). Moreover, the translated introns tended to be more positively charged (Fig. 2D) than CDS and upstream exons. This agrees with established traits of IDRs ^47–49^. The variance in hydrophobicity was marginal (Fig. 2E). We note that similar trends were observed irrespective of analysis window sizes (Extended Data Fig. 2B-D).

The amino acids serine (S), proline (P), glycine (G), and arginine (R) were highly overrepresented in translated intron regions (Fig. 2F). We observed this enrichment even when changing the reading frame by -1 or +1 nucleotides (Extended Data Fig. 2E). As P, G, and R are encoded by multiple numbers of GC rich codons (*e.g.*, CCN for P, GGN for Gly, and CGN for R), we analyzed the GC content of the introns throughout the length. As expected ^50, 51^, the GC content is higher towards the 5′ end of the intron, thereby increasing the likelihood of encountering S, P, G, or R codons and therefore producing stretches of low complexity in the resulting proteins (Extended Data Fig. 2F). The presence of low complexity regions (LCRs) might lead to phase separation and/or to prion-like aggregation of the chimeric protein products ^52–55^.

### A subset of intron-derived peptides are condensation-prone

The enrichment of IDRs in intron-derived peptides prompted us to test the condensation propensity of the chimeric proteins. For this purpose, we performed BONCAT proteomic analysis of the cellular soluble and insoluble fractions. Regardless of the limited number of detected intron-derived peptides, we observed that the chimeric peptides detected in the pellet of SSA-treated cells appeared more prevalent than in the supernatant (Fig. 3A). The protein ferritin heavy chain 1 (FTH1) represents a remarkable example, as ribosome profiling indicated intron translation (Fig. 3B) and its truncated form was highly enriched in the pellet fraction of the BONCAT experiments (Fig. 3A, left panel). The translated intron of FTH1* possessed an LCR and showed a propensity for disorder (Fig. 3C). Similar LCRs were found in other condensate-prone, translated introns (RAB5IF* and ABHD11*) (Fig. 3A, right panel and Extended Data Fig. 3A).

**Fig. 3:**
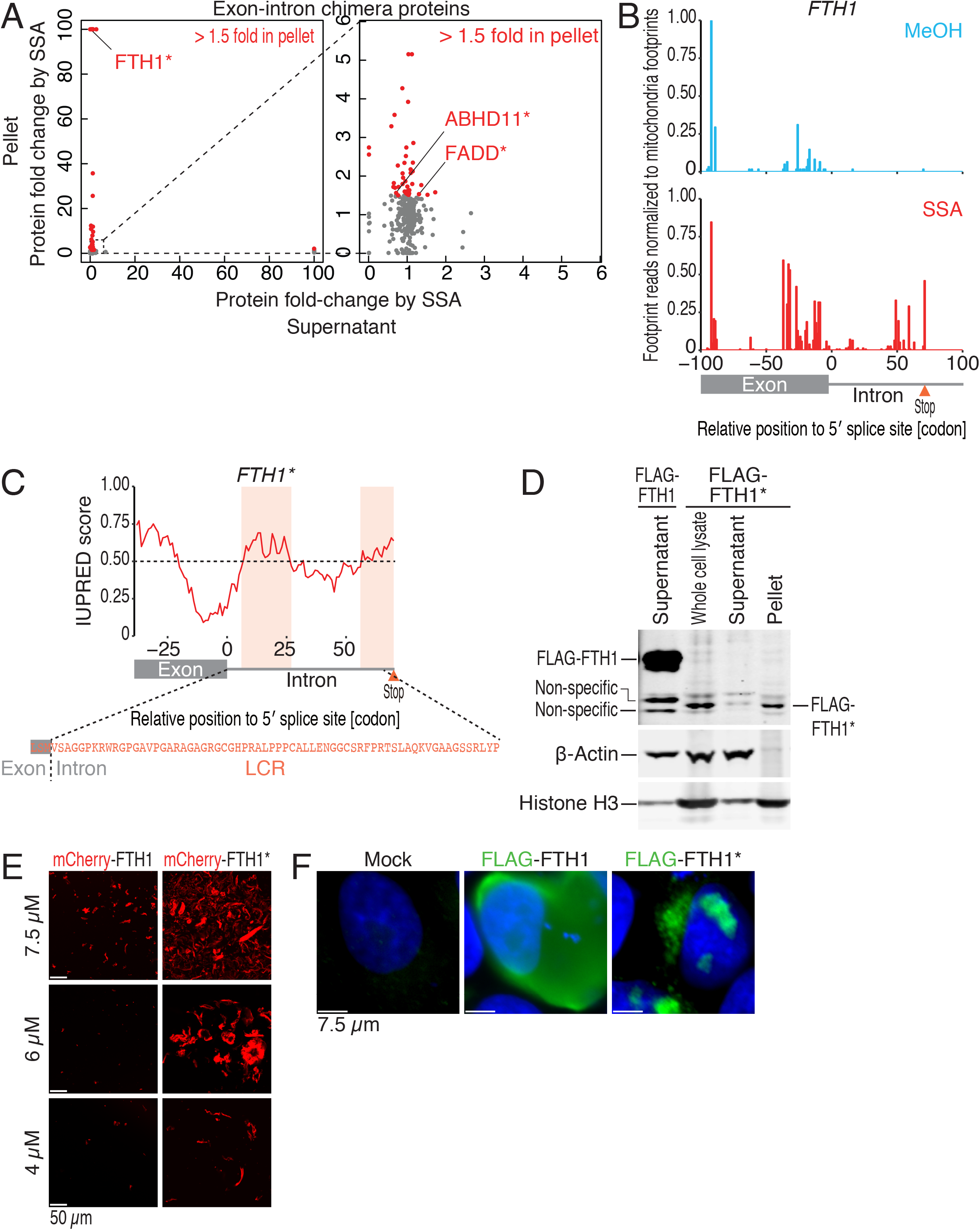
A subset of intron-derived peptides is condensation-prone. (A) Exon-intron chimeric proteins from the centrifuge supernatant (horizontal axis) and pellet (vertical axis) of HeLa S3 cells treated with 100 ng/ml SSA for 6 h were quantified by BONCAT. The right graph is zoomed-in graph of the left. Peptides originating from the entire exon-intron region were considered. Red: exon-intron chimera proteins enriched 1.5-fold or more in pellet fraction. (B) The accumulation of ribosome footprints on *FTH1* introns under splicing perturbation. Reads were normalized to the sum of mitochondrial footprint reads. (C) Prediction of disordered regions in FTH1* by IUPred2A ^46^. The LCR sequence predicted by SEG ^91^ is shown. (D) Western blot of FLAG-tagged FTH1* ectopically expressed in HeLa S3 cells. The cell lysate was further fractionated by centrifugation. (E) Confocal micrographs of recombinant FTH1 and FTH1* proteins fused to mCherry. Scale bars are 50 µm. (F) HeLa S3 cells were transfected with either mock, FLAG-FTH1, or FLAG-FTH1* vectors for 24 h. Representative fluorescence micrographs show the distribution of FLAG-tagged protein, immunostained (green). Nuclei were stained with Hoechst dye (blue). Scale bars are 7.5 µm.

The condensate-prone character of the ectopically expressed truncated FTH1* was validated through fractionation by centrifugation followed by Western blotting (Fig. 3D). Similar condensates were also obtained for FADD* and IRF2BP2* (Extended Data Fig. 3B) when they were expressed in cells. Although RAB32* was not supported by BONCAT, we did find a likely condensate prone feature in this protein (Extended Data Fig. 3C) when seeking candidate sequences in our ribosome profiling data (Extended Data Fig. 3D). Our newly identified truncated proteins were distinct from the previously reported p27*, as it constitutes a shortened, but soluble and functional polypeptide (Extended Data Fig. 3E).

In addition, the recombinant protein FTH1*, but not FTH1, formed aggregates by self-associated *in vitro* in a concentration-dependent manner (Fig. 3E). Similar FTH1* condensates were found in cells as well, when it was expressed as FLAG-tagged protein (Fig. 3F).

### Condensation-prone intron-derived peptides are proteotoxic

Given that translation of improperly spliced transcripts leads to the production of truncated and cellular condensates/aggregates (Fig. 3), it appears feasible that these imperfect peptides produced upon SSA trigger a proteotoxic stress response. To test this idea, we investigated whether c-Jun N-terminal kinase (JNK), a multifaceted kinase that responds to different cues, including proteotoxic stress ^56, 57^, is activated upon SSA treatment. Consistent with this scenario, we detected that treatment with SSA (Fig. 4A) induced the activation of JNK signaling, as seen by its increased JNK phosphorylation, similar to the level induced by proteasome inhibitor MG132 (Fig. 4A). Whereas JNK activation was not detected when acetyl-SSA (Ac-SSA) (Extended Data Fig. 4A), an inactive SSA derivative ^9^ was used, it did occur with another SF3B1 inhibitor, pladienolide B (PlaB) ^19, 58, 59^ (Extended Data Fig. 4B), which also induces intron retention (Extended Data Fig. 4C). We could therefore rule out off-target effects and conclude that splicing inhibition causes proteotoxic stress.

**Fig. 4:**
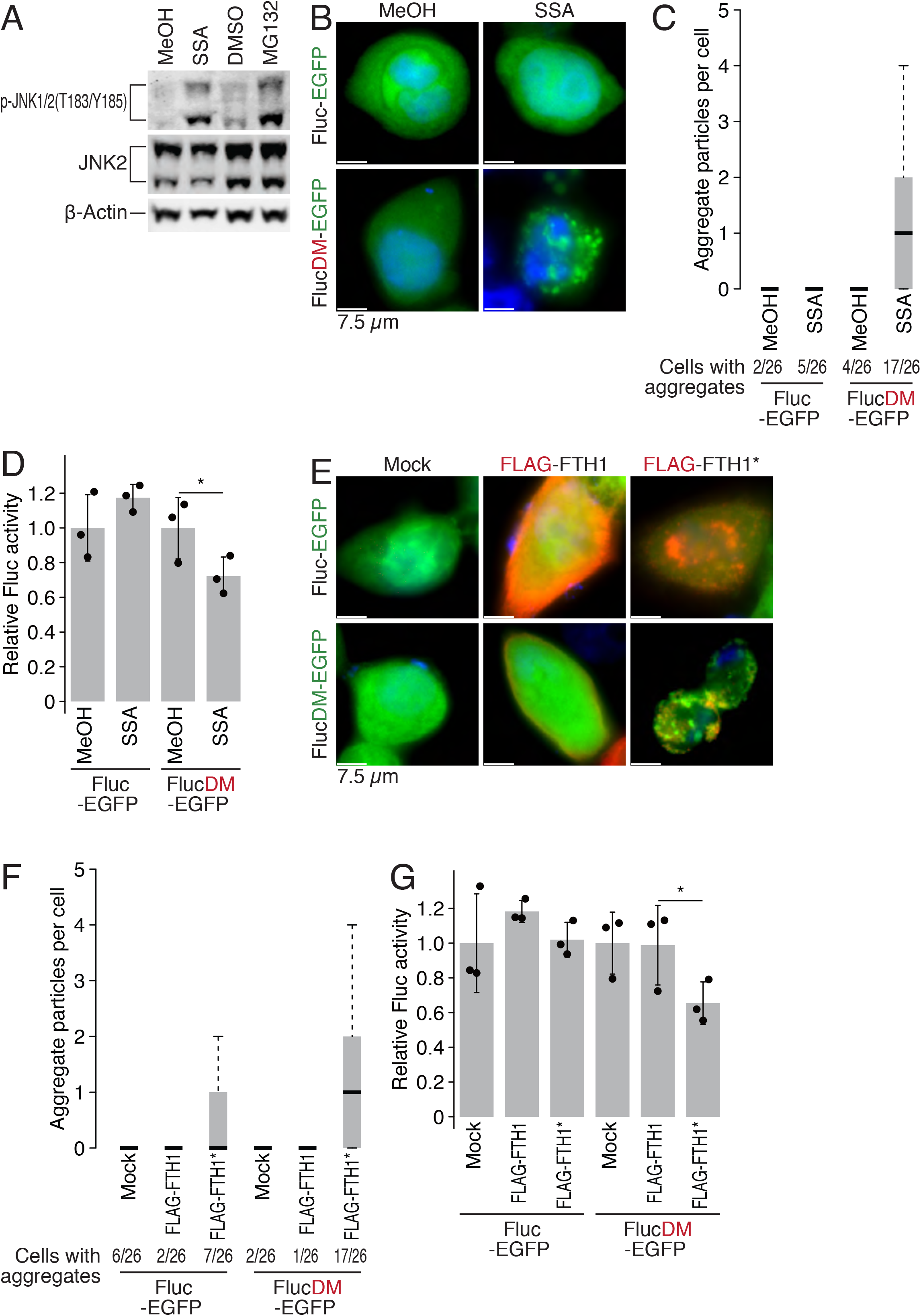
Condensation-prone intron-derived peptides are proteotoxic. (A) HeLa S3 cells were either treated with 100 ng/ml SSA or 0.5 µM MG132 for 10 h, and the cell lysates were immunoblotted with the indicated antibodies. (B) HeLa S3 cells were transfected for 48 h with a Fluc-based sensor reporter construct encoding Fluc-EGFP or FlucDM-EGFP and subsequently incubated for 10 h with either 100 ng/ml SSA or solvent MeOH. Representative fluorescence micrographs show the distribution of EGFP signals (green) in cells. Nuclei were stained with Hoechst dye (blue). Scale bars are 7.5 µm. (C) The number of aggregated GFP per cell in (B). A minimum of 26 cells were quantified in each condition. (D) Firefly luciferase activities in (B). Data were normalized to the total GFP protein detected by Western blot (MeOH control values were set to 1). Data represent the mean and s.d. (n = 3). *P* values were calculated using Student’s *t*-test (one-tailed). *, *P* < 0.05. (E) HeLa S3 cells were cotransfected with the Fluc-based sensor reporter construct encoding Fluc-EGFP or FlucDM-EGFP and either mock, FLAG-FTH1, or FLAG-FTH1* vectors for 48 h. Representative fluorescence micrographs show the distribution of EGFP signal (green) in cells. FLAG-tagged protein was immunostained (red). Nuclei were stained with Hoechst dye (blue). Scale bars are 7.5 µm. (F) The number of aggregated GFP per cell in (E). A minimum of 26 cells were quantified in each condition. (G) Firefly luciferase activities in (E). Data were normalized to the total GFP protein detected by Western blot (MeOH control values were set to 1). Data represent the mean and s.d. (n = 3). *P* values were calculated using Student’s *t*-test (one-tailed). *, *P* < 0.05.

To visualize the proteotoxic stress response in individual cells, we used a firefly luciferase (Fluc) reporter and its conformationally unstable version (R188Q-R261Q double mutant or DM), which requires chaperone surveillance to fold, fused to GFP to assess a possible imbalance in proteostasis ^60^. Under proteotoxic stress, chaperones become limiting, decreasing the solubility of the sensor protein and thereby preventing the luminescent reporter from proper protein folding. As expected, SSA treatment of cells harboring the FlucDM-EGFP reporter produced GFP aggregates (Fig. 4B and 4C), as well as reduced luciferase activity (Fig. 4D). These results corroborate the suspected proteostasis imbalance induced by splicing modulation. While proteotoxic stress under SSA treatment is likely the cumulative result of many truncated peptides, we investigated whether the ectopic expression of individual truncated proteins including their intron-derived sequences recapitulates the phenotype of SSA treatment at least in part. The expression of condensate-prone FTH1* induced an imbalance in proteostasis, as observed by condensate formation (Fig. 4E and 4F), along with the reduction of firefly luciferase activity (Fig. 4G), in the proteostasis reporter.

Our results suggest that an overload of toxic, condensate-prone proteins generated by splicing perturbation elicits proteotoxic stress, leading to activation of the stress-activated protein kinase JNK.

### SSA induces global translation inhibition

During our analysis of ribosome profiling, we noticed that the impact of SSA was not restricted to intron translation. With the reported decrease in transcription and mRNA export upon SSA treatment ^9, 37, 38^, some concomitant decrease in protein synthesis would appear natural. Measuring the absolute change in translation by ribosome profiling using mitochondrial ribosome footprints as an internal control (see Materials and Methods for details), we found that splicing inhibition led to a far more drastic decrease (fourfold) in translation than could be explained by a decreased number of transcripts alone (twofold) (Fig. 5A). Metabolic labeling of nascent peptides using *O*-propargyl-puromycin (OP-puro) followed by fluorophore conjugation ^61, 62^ further corroborated a strong inhibition of protein synthesis in the presence of SSA (Fig. 5B and Extended Data Fig. 5A).

**Fig. 5:**
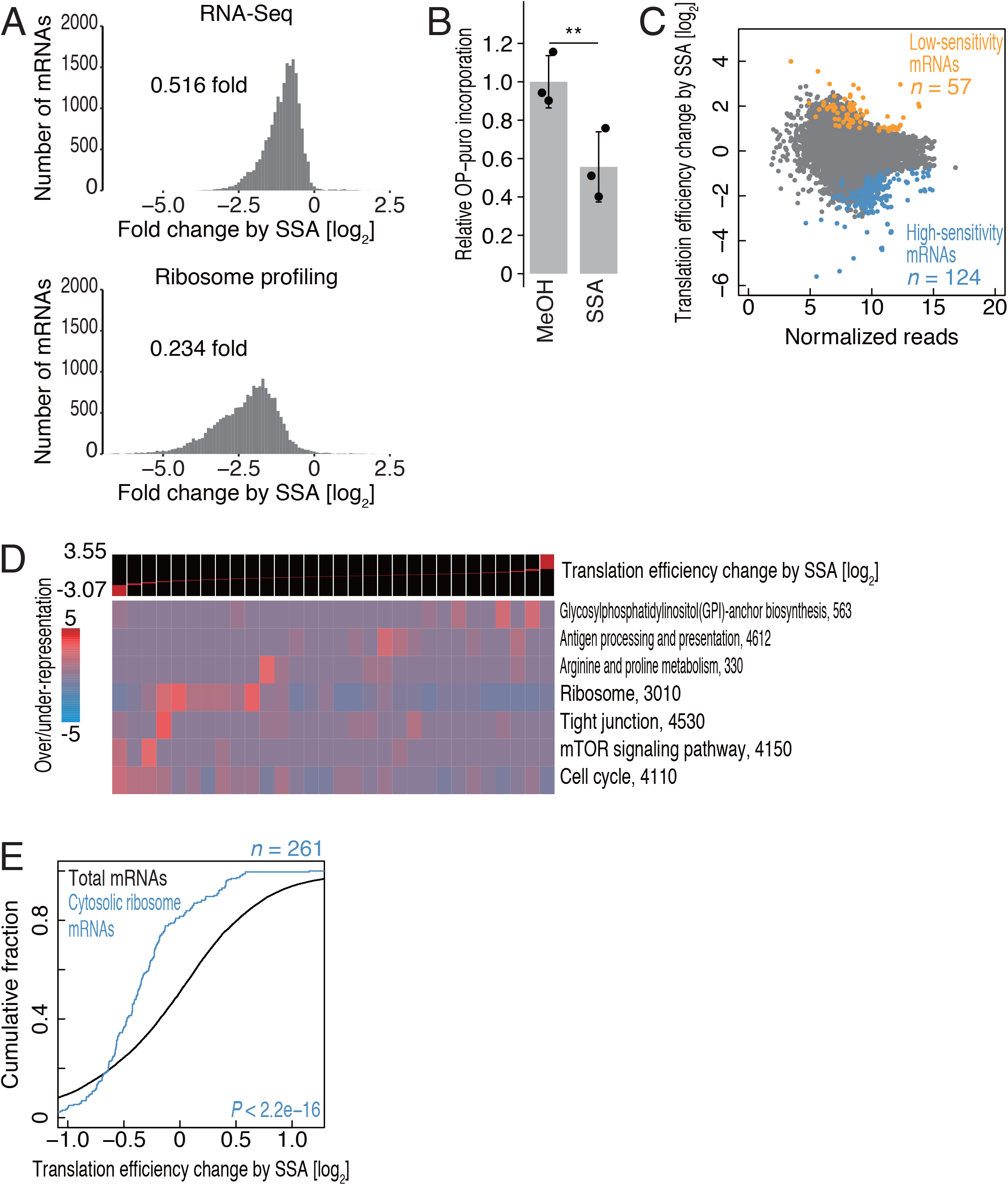
Global translation was inhibited upon splicing modulation. (A) Histograms showing absolute change in RNA-Seq reads (upper panel) and ribosomal footprints (lower panel) under SSA treatment. Data were normalized to the total number of reads from mitochondrial genome-encoded transcripts used as internal spike-ins. A median-fold change is shown. The bin width is 0.1. (B) Bulk translation change upon SSA treatment in HeLa S3 cells was monitored by OP-puro. Nascent peptides with incorporated OP-puro were visualized by click reaction with azide-conjugated IR-800 dye and quantified. Data represent the mean and s.d. (n = 3). *P* values were calculated using Student’s *t*-test (one-tailed). **, *P* < 0.01. (C) MA plot of translation efficiency change during SSA treatment plotted against normalized RNA-Seq reads. FDR < 0.05 was set for the definition of low-sensitivity mRNAs and high-sensitivity mRNAs. (D) KEGG pathway analysis based on the differential change in translation efficiency, visualized by iPAGE ^89^. (E) Cumulative distribution of cytosolic ribosome mRNAs in translation efficiency change during SSA treatment. The significance was calculated by Wilcoxon’s test.

We explored whether differential changes in translation efficiency of individual mRNAs would have functional implications. Translation efficiency, the ratio between footprint and mRNA-sequencing counts, presents a potent means to quantify actually occurring translation, as it takes transcript abundance into account. We observed differential changes across a number of transcripts during SSA treatment (Fig. 5C). Based on the Kyoto Encyclopedia of Genes and Genomes (KEGG), we found that the translation efficiencies of ribosomal proteins were particularly reduced (Fig. 5D). Similar results were obtained through Gene Ontology (GO) analysis (Extended Data Fig. 5B). The downregulation of translation efficiency in mRNAs coding for components of the cytosolic ribosome stood out from the other affected genes (Fig. 5E).

### SSA-mediated JNK activation leads to mTORC1 inhibition

The translation decrease from ribosomal protein genes is a hallmark of mTORC1 inhibition ^63, 64^. Next, we tested the possibility that SSA leads to mTORC1 inactivation, as mTORC1 can be deactivated upon sensing proteotoxic stress via JNK (Extended Data Fig. 6A) ^57^. mTORC1 — a complex consisting of mTOR kinase at its core and the accessory proteins DEPTOR, RAPTOR, PRAS40, and mLST8 — is the master regulator of protein synthesis ^65, 66^ through direct phosphorylation of several translational key regulators, such as eukaryotic translation initiation factor 4E (eIF4E) binding protein 1 (4EBP1) and ribosomal protein S6 kinase beta-1 (S6K1).

**Fig. 6:**
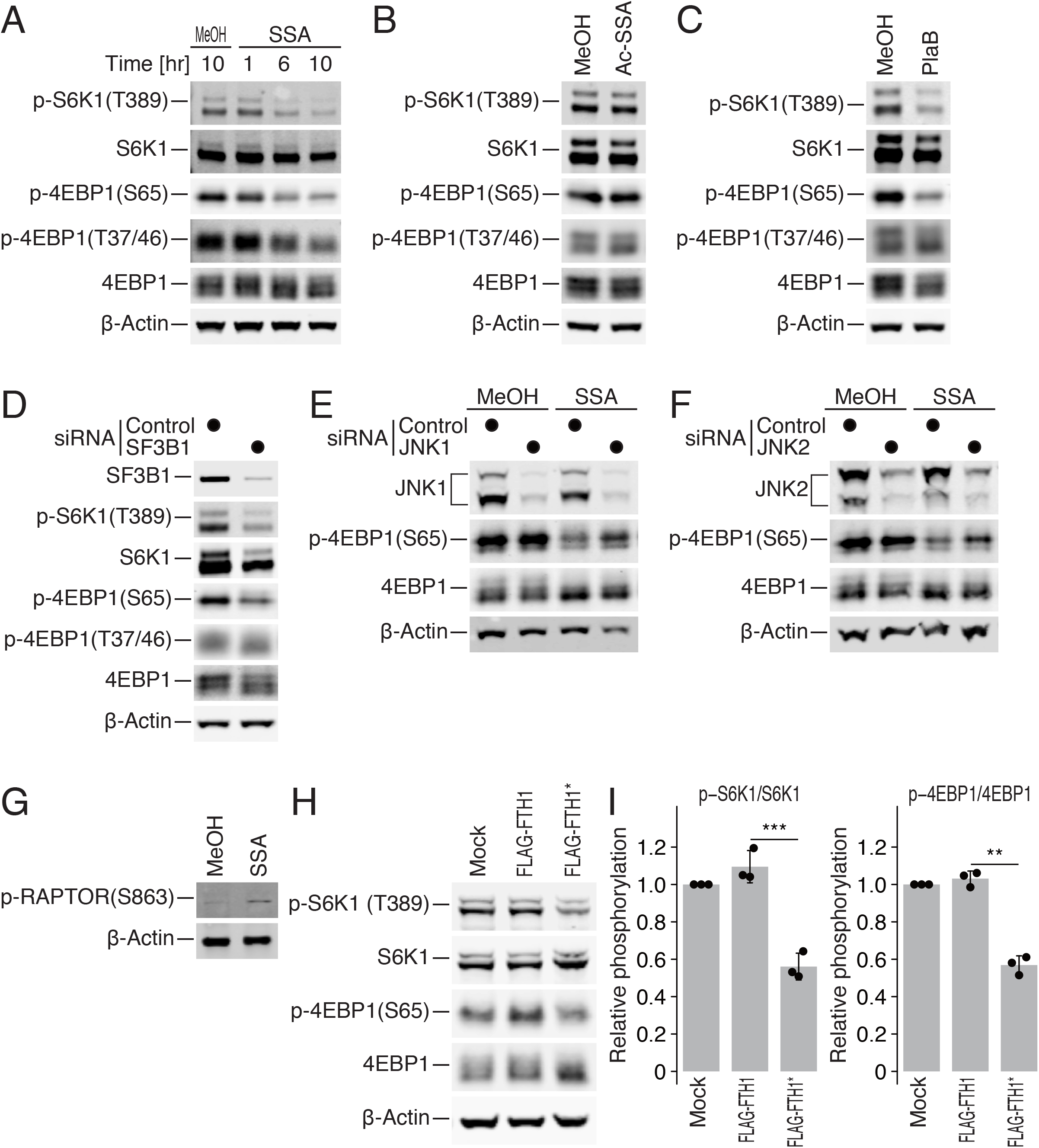
SSA-mediated JNK activation leads to mTORC1 inhibition. (A) Western blot for mTORC1 substrates S6K1 and 4EBP1 and their phosphorylated forms. HeLa S3 cells were either treated with MeOH solvent or 100 ng/ml SSA for different time periods. (B and C) Western blot of the indicated proteins from HeLa S3 cells treated with 100 ng/ml Ac-SSA (B) and 1 µg/ml PlaB (C) for 10 h. (D) HeLa S3 cells were either transfected with control siRNA or siRNA targeting SF3B1 mRNA for 36 h. Cell lysates were immunoblotted with the indicated antibodies. (E and F) JNK1 (E) or JNK2 (F) was knocked down in HeLa S3 cells before treatment with 100 ng/ml SSA for 10 h. The cell lysates were immunoblotted with the indicated antibodies. (G) HeLa S3 cells were either treated with 100 ng/ml SSA or MeOH solvent for 10 h, and the cell lysates were immunoblotted with the indicated antibodies. (H and I) Representative Western blots for either phosphorylated or bulk S6K1 and 4EBP1 in stable HEK 293 cells, which express FLAG-FTH1 or FLAG-FTH1*, are shown (H) and quantified (I). The data represent the mean and s.d. (n = 3). *P* values were calculated using Student’s *t*-test (one-tailed). **, *P* < 0.01; ***, *P* < 0.001.

To test for mTORC1 inhibition by SSA, we examined the phosphorylation status of mTORC1 substrates 4EBP1 (at Thr37/46 and Ser65) and S6K1 (at Thr389). We observed dephosphorylation of these proteins in the presence of SSA (Fig. 6A and Extended Data Fig. 6B and 6C). In contrast, Ac-SSA did not cause any dephosphorylation of mTORC1 substrates (Fig. 6B).

mTORC1 inhibition was not specific to SSA but could be reproduced when inhibiting the SF3B1 complex by other means. We repeated the above assays with PlaB and knockdown of SF3B1. Either method also induced intron retention (Extended Data Fig. 4C and 6D), and both conditions recapitulated the dephosphorylation of 4EBP1 and S6K1 (Fig. 6C and 6D). Although this provided strong evidence that the observed effect on mTORC1 signaling must depend on aberrant splicing, we used an *in vitro* assay with recombinant mTORC1 protein to confirm that neither PlaB nor SSA directly inhibited its kinase activity (Extended Data Fig. 6E).

We assessed whether the observed JNK activity was responsible for mTORC1 inactivation. Individual knockdown of JNK variants JNK1 and JNK2 recovered 4EBP1 phosphorylation in the presence of SSA (Fig. 6E and 6F).

Proteotoxic stress-activated JNK mediates the disassembly of the mTORC1 complex via phosphorylation of mTORC1 component RAPTOR on S863 (Extended Data Fig. 6A) ^57^. We observed increased phosphorylation of RAPTOR S863 in the presence of SSA (Fig. 6G), supporting a functional link between JNK activation and mTORC1 inhibition.

JNK-mediated mTORC1 inhibition most likely originates from SSA-induced truncated peptide-derived condensates that cause proteotoxic stress. In line with our experiments on proteotoxic stress (Fig. 4D-F), we observed that the expression of condensate-prone FTH1* inhibited mTORC1, as demonstrated by the dephosphorylation of 4EBP1 and S6K1 (Fig. 6H and I). In sum, our results suggest that an overload of toxic, condensate-prone proteins generated by splicing perturbation leads to mTORC1 deactivation via activation of JNK.

### SSA mimics translation repression by mTORC1 inactivation

Given the correspondence between 4EBP1 dephosphorylation (Fig. 6) and translation repression (Fig. 5), we inferred that SSA reduces protein synthesis through mTORC1 inactivation. To test this hypothesis, we directly compared the translation change evoked by SSA and ATP-competitive mTOR inhibitor pp242. Based on ribosome profiling data, transcripts sensitive to pp242^67^ proved also sensitive to SSA (Fig. 7A).

**Fig. 7:**
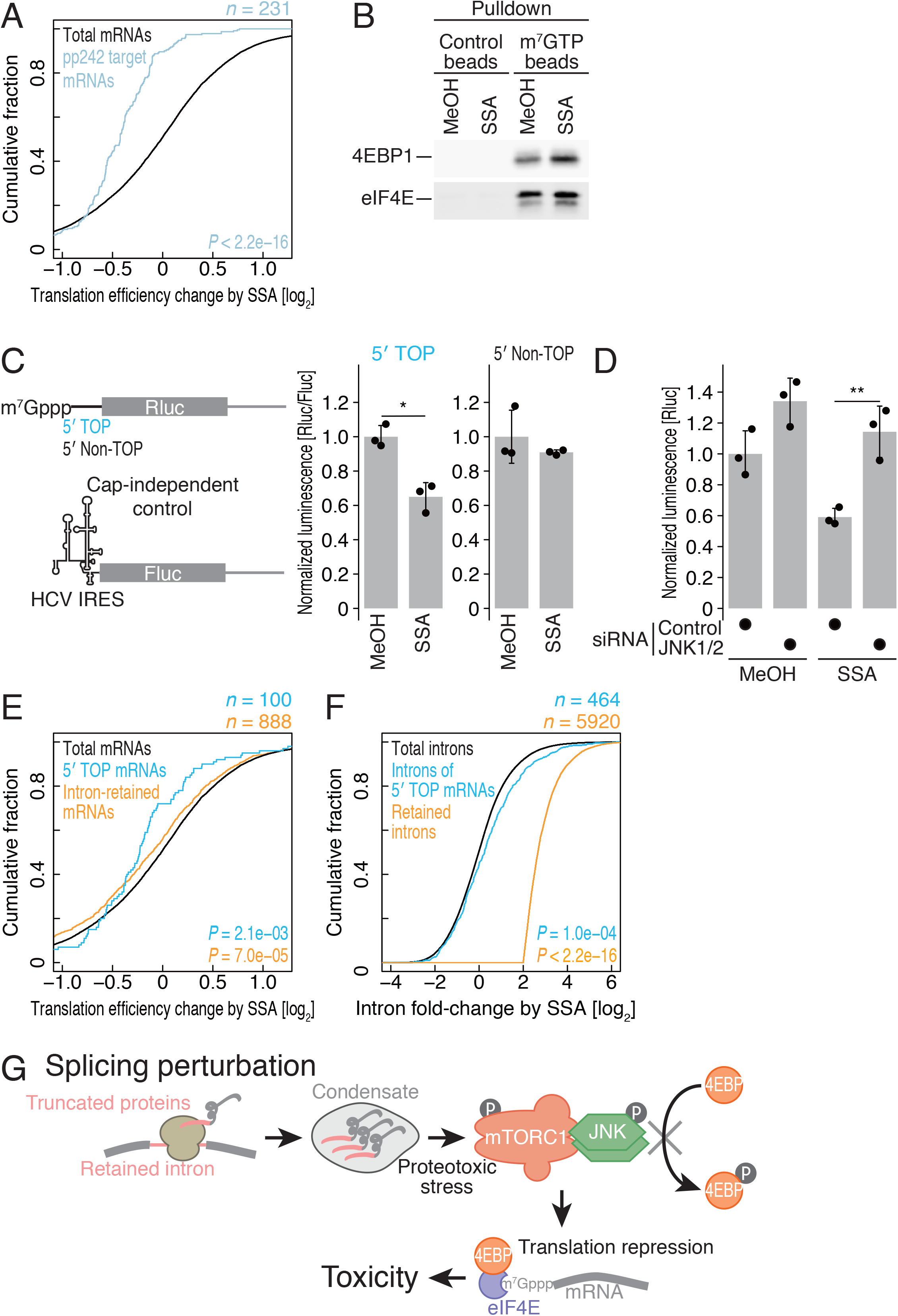
Characterization of translation repression induced by SSA. (A) Cumulative distribution of pp242-sensitive mRNAs ^67^ in translation efficiency change during SSA treatment. Significance was calculated by Wilcoxon’s test. (B) Proteins copurified with m^7^G-cap beads were used for Western blotting. HeLa S3 cells were either treated with the MeOH control or SSA for 10 h. (C) Reporter *Renilla* luciferase mRNA fused downstream to the TOP motif containing the 5′ UTR of *EIF2S3* or the non-TOP 5′ UTR of *ATP5O* (Extended Data Fig. 7A) and firefly luciferase mRNAs fused downstream to the HCV IRES were transfected into HeLa S3 cells for 4 h after 2 h of treatment with 100 ng/ml SSA. Data represent the mean and s.d. (n = 3). *P* values were calculated using Student’s *t*-test (one-tailed). *, *P* < 0.05. (D) 5′ TOP reporter assay in HeLa S3 cells with 100 ng/ml SSA. Cells were transfected with siRNAs against JNK1 and 2. Data represent the mean and s.d. (n = 3). *P* values were calculated using Student’s *t*-test (one-tailed). **, *P* < 0.01. (E and F) Cumulative distribution of 5′ TOP mRNAs (curated from defined 5’ UTR ^90^) and intron-retained mRNAs in translation efficiency change during SSA treatment (E) and in intron read change during SSA treatment (F). The significance was calculated by Wilcoxon’s test. (G) Model for translation attenuation upon splicing modulation. Under splicing attenuation, introns of pre-mRNAs are retained. A subset of RNAs is transported into the cytoplasm and is translated. Synthesized proteins from intron-retained RNA possessing a condensate-prone character lead to proteotoxic stress and JNK-mediated phosphorylation of mTORC1 components. Inhibition of mTORC1 results in reduced phosphorylation of its target proteins, including 4EBP and S6K1, and a subsequent decrease in protein biosynthesis for tumor toxicity.

Since dephosphorylated 4EBP1 binds to the cap-binding protein eIF4E, thereby inhibiting eIF4F (a complex of eIF4E, G, and A) formation ^68^, we tested whether SSA inhibits cap-dependent translation. Consistent with the dephosphorylation of 4EBP1 upon SSA treatment (Fig. 5), pulldown experiments using cell lysates on m^7^G-cap beads showed stronger association between 4EBP and eIF4E in SSA-treated cells (Fig. 7B).

Given that mRNAs coding for ribosomal proteins frequently possess a terminal oligopyrimidine motif (5’ TOP) motif in their 5’ untranslated regions (UTRs), which particularly sensitizes their translation to mTORC1 inhibition ^69^, we investigated their translation efficiency under SSA. To test the sensitivity of 5’ TOP motif-containing mRNAs to SSA-induced translation repression, irrespective of their splicing status, we transfected *in vitro* synthesized intronless *Renilla* luciferase reporters into HeLa S3 cells. Firefly luciferase under the control of the HCV-internal ribosome entry site (IRES), which allows translation initiation independent of the eIF4F complex, served as control ^70^ (Fig. 7C). In agreement with mTORC1 inactivation, SSA led to strongly reduced expression of 5’ TOP motif-containing mRNAs (Fig. 7C) compared to non-TOP controls (Extended Data Fig. 7A). Furthermore, the SSA-induced reduction in 5’ TOP reporter translation could be reversed when JNK1 and JNK2 were knocked down (Fig. 7D and Extended Data Fig. 7B). Thus, our controlled reporter assay corroborated selective inhibition of mTORC1 by splicing modulation through activation of JNK.

These observations could not be explained by the action of SSA on splicing alone. The number of ribosomal footprints on intron-retained mRNAs will naturally decrease, as translation will halt at the first intronic stop codon, effectively decreasing the space available to ribosomes. Consistent with this scenario, we observed a reduction in the translation efficiency of the intron-retained mRNAs upon SSA treatment (Fig. 7E). However, the change in the translation efficiency of the 5’ TOP mRNA was more prominent (Fig. 7E). Conversely, 5’ TOP mRNA hardly contained any retained introns even in the presence of SSA (Fig. 7F). Therefore, these data demonstrate that selective inhibition of mTORC1-mediated translation upon SSA treatment, but not splicing defects *per se*, accounts for the observed decrease in protein output.

Our results indicate that toxic, condensate-prone proteins generated by splicing perturbation leads to mTORC1-dependent translational repression via JNK activation (Fig. 7G).

## Discussion

The function of peptides generated outside of the coding regions have long been overlooked. Recent reports shed light on the role of noncanonical peptides. Substantial numbers of functional noncanonical human micropeptides have been reported from 5′ UTRs (or upstream ORFs, uORFs) playing an essential physiological role ^71^. Amino acid sequences encoded in the 3′ UTRs were reported to suppress C-terminally extended proteins generated by stop codon readthrough ^72^. Along with these studies, the functional exon-intron chimeric protein related in this study expands the cellular proteome.

The importance, benefits, and burdens of introns remain a topic of discussion ^73^. Even in intron-poor budding yeast, with only 300 introns across its entire genome, these intervening sequences bear important cellular functions, such as sensing and mediating the starvation response ^74, 75^. In the stationary phase of yeast, decreased TORC1 activity leads to intron stabilization, subsequent spliceosome sequestration, and attenuation of protein expression from intron-containing mRNAs such as ribosomal proteins. Although our study in mammals shows the opposite regulatory direction — intron retention triggering mTORC1 inhibition, the key molecular complex (TORC1) and regulatory targets (translation machinery) are the same. This may indicate that the basic properties of introns are conserved between yeast and humans.

The condensate-prone feature of low complexity peptides generated from intronic RNA sequences highlights the hidden function of exon-intron chimeric proteins. The high GC content around the 5′ splice site and its role in splicing has been found by earlier studies ^50, 51^. Our study suggested the alternative role to encode condensation-prone P-, G-, and R-rich region. Cellular condensates in the form of liquid, gel, and glass states have gained great attention since they have roles in concentrating a set of molecules to enhance biochemical reaction rates, sequestering complexes in response to environmental stimuli, compartmentalizing cellular space as membrane-less organelles, and signaling switches ^52–55^.

Although we demonstrated that FTH1* condensates function as a proteotoxic signal inducer, other translated introns with low complexity regions may carry alternative functions.

The mechanism by which splicing modulators exhibit their antitumor activity has remained speculative. Although the direct alteration of splicing in key genes offers a straightforward explanation, it is limited to a small number of cancer cells. For example, in a subset of cancer cells that depend on high levels of Mcl-1 — a Bcl2 family apoptosis regulator — for survival, splicing modulators induce cell death by splice variant switching from the antiapoptotic Mcl-1L to the proapoptotic Mcl-1S isoform ^24–26^. Splicing modulators are particularly potent in inhibiting many hematological malignancies harboring splicing factor mutations, which supports the idea of synthetic lethality ^16, 21, 22^. Similarly, Myc-activated tumor cells are vulnerable to splicing modulation due to a high demand of spliceosomal activity ^23^. Additionally, splicing modulation drastically changes the transcription of particular genes: VEGF gene expression is downregulated ^27–29^, while NF-kB-dependent transcription is activated ^76^. In this study we present a more general mechanism, by which mTORC1 inhibition represses protein synthesis in response to splicing inhibition. As increased translation through hyperactive mTORC1 has been a hallmark of many human cancers, it is plausible that suppression of mTORC1 activity contributes to the anticancer activity of splicing modulators. mTOR inhibitors inhibit the expression of HIF-1 dependent genes including VEGF and activate apoptotic pathways in cells defective in tumor suppressors such as p53 and PTEN ^65, 77^. Indeed, mTOR inhibitors temsirolimus and everolimus have already been used for cancer treatment ^65, 77^.

This scenario is not mutually exclusive with other possibilities. The downregulation of tumorigenic genes through reduced protein output could form the basis of antitumor activity. We observed strong translational repression of cancer-related genes such as *CYR61*, *PABPC4*, *CDK9*, *SGK1*, *KIF23*, and *RAE1* under SSA treatment (Extended Data Fig. 7C). It is further possible that some other truncated proteins gain functions as “neopeptides” to induce cell death, cell senescence or differentiation. More interestingly, the prospect of SSA-induced exon-intron chimeric proteins may generate new antigens, which could form the basis for future cancer immunotherapy ^48, 78, 79^. Antigen processing and presentation genes appeared upregulated in translation efficiency under SSA, possibly reflecting the production of the neoantigens (Fig. 5D). Multiple mechanisms, including downregulation of mTORC1, may cooperate towards potent antitumorigenic activity when splicing is inhibited. Our genome-wide transcriptome/translatome data provide a useful resource for further investigation.

Although we chemically induced intron retention, the widespread presence of IDR-encoded introns suggests genuine physiological roles in contexts where intron retention is induced, such as in cancer ^6^, aging ^80^, and heat shock ^81^. Under these conditions, mTORC1 inhibition evoked by cellular condensates may feed back into translation and mitigate the synthesis of harmful proteins.

## Acknowledgments

Rei Yoshimoto, Daisuke Kaida, and all the members of the Iwasaki and Yoshida laboratories for constructive discussions, technical help, and critical reading of the manuscript and the Support Unit for Bio-Material Analysis, RIKEN CBS Research Resources Division for technical help. PlaB was kindly provided by Eisai Co., Ltd. pCI-neo Fluc-EGFP and pCI-neo FlucDM-EGFP were kind gifts from Franz-Ulrich Hartl. M.Y. was supported in part by the Grant-in-Aid for Scientific Research (S) (JP19H05640) from the Japan Society for the Promotion of Science (JSPS) and the Grant-in-Aid for Scientific Research on Innovative Areas “Ubiquitin new frontier driven by Chemo-technologies” (JP18H05503) from the Ministry of Education, Culture, Sports, Science and Technology (MEXT). S.I. was supported by the Grant-in-Aid for Scientific Research on Innovative Areas “nascent chain biology” (JP17H05679) from MEXT, the Grant-in-Aid for Young Scientists (A) (JP17H04998), Challenging Research (Exploratory) (JP19K22406), and the Grant-in-Aid for Transformative Research Areas (B) (JP20H05784) from JSPS, AMED-CREST, AMED (JP20gm1410001), the Pioneering Projects (“Cellular Evolution”) and the Aging Project from RIKEN, and the Takeda Science Foundation. J.K.C.S. was supported by a Grant-in-Aid for Early-Career Scientists (JP20K15420) from JSPS. DNA libraries were sequenced by the Vincent J. Coates Genomics Sequencing Laboratory at UC Berkeley, supported by the NIH S10 OD018174 Instrumentation Grant. Computations were supported by Manabu Ishii, Itoshi Nikaido, the Bioinformatics Analysis Environment Service on RIKEN Cloud, and supercomputer HOKUSAI SailingShip in RIKEN ACCC. J.K.C.S. was a recipient of the Japanese Government (MEXT) Scholarship.

## Author Contributions

J.K.C.S. performed the experiments and bioinformatics analyses with the help of M.M. and K.K.; S.I., T.S.P., and M.Y. supervised the overall experiments; T.S. and N.D. conducted mass spectrometry; J.K.C.S., T.S.P., and S.I. wrote the manuscript with the inputs from all the authors.

## Materials and Methods

### Cell culture

HeLa S3 and HEK293 T-REx (Thermo Fisher Scientific) cells were maintained in DMEM supplemented with 10% FBS in a humidified incubator at 5% CO_2_ and 37°C. The stable cell lines of HEK293 T-REx were selected following the manufacturer’s instructions.

### Compounds

SSA and PlaB were dissolved in methanol (MeOH). MG132 (Wako chemicals), rapamycin (Wako chemicals), and pp242 (Sigma-Aldrich) were dissolved in DMSO.

### Ribosome profiling and RNA-Seq

HeLa S3 cells were grown in 10-cm dishes at 70-80% confluency and treated with SSA (100 ng/ml) or its solvent MeOH for 6 h before lysis. The libraries for ribosome profiling were prepared as described earlier ^67, 82^ and sequenced by a HiSeq4000 sequencer (Illumina).

Total RNA was extracted from the same lysate used for ribosome profiling with TRIzol LS (Thermo Fisher Scientific) and Direct-zol RNA MicroPrep Kits (Zymo Research). The libraries were prepared with the TruSeq Stranded mRNA Library Prep Kit followed by rRNA removal with Ribo-Zero Gold (Illumina) and were sequenced on a HiSeq4000 sequencer (Illumina).

### Data analysis

#### Ribosome profiling

First, the 3′ adaptor sequence (5′-AGATCGGAAGAGCACACGTCTGAA-3′) was trimmed using fastx clipper (Hannon lab, http://hannonlab.cshl.edu/fastx_toolkit/commandline.html) and then demultiplexed according to the barcode sequences in the linkers. We used Bowtie2 ^83, 84^ to map the clipped reads to a human rRNA, tRNA, snoRNA, snRNA, and microRNA reference database and captured unaligned reads. These reads were mapped to the human genome [hg19; known reference genes from University of California, Santa Cruz (UCSC)] using Tophat ^85^. The PCR duplicates were eliminated thanks to a randomized sequence stretch in the linkers. Empirical estimation of nucleotides on the ribosomal A-site was performed on the basis of footprint length: 15 for 27-28 nt reads and 16 for 29-31 nt reads. Translated introns were defined as follows: (i) intron length > 200 nt, (ii) at least one footprint on the intron, and (iii) 5 or more footprints on CDS exons. Except for the first and last 5 amino acids, the reads from CDS were counted. The relative enrichment of reads was calculated by DESeq ^86^.

We measured the change in overall translation by normalizing the number of cytosolic ribosome footprints to those from mitochondrial ribosomes ^67^, as SSA treatment does not appear to impact mitochondrial protein synthesis.

#### RNA-Seq

After clipping the adaptor sequence (5′-AGATCGGAAGAGCACACGTCTGAACTCCAGTCAC-3′), the reads were processed as described above for ribosome profiling data analysis.

Mixture-of-isoforms (MISO) ^87^ was used to assess alternative splicing events upon splice inhibition across a database of annotated splice events (https://miso.readthedocs.io/en/fastmiso/) using the following filtering criteria: (i) both inclusion and exclusion reads were ≥ 1 such that (ii) the sum of inclusion and exclusion reads was ≥ 10, (iii) the absolute values of the difference for “Percent Spliced In” (ΔPSI) between vehicle and SSA were ≥ 0.2, and (iv) the Bayes factor was ≥ 10.

For differential intron retention analysis, reads from introns were analyzed by DESeq ^86^. The reads were processed as follows: (i) the number of reads was ≥ 5, (ii) introns within the ORF were considered, (iii) MAXENT score, strength of 5’ splice sites ^88^ was > 2.5 to minimize the background, and (iv) corresponding mRNAs had ≥ 5 reads in the ORFs. Retained introns were defined as FDR < 0.01 and read enrichment ≥ 4-fold.

#### Translation efficiency

The differential change in ribosome footprint reads over RNA-Seq reads was calculated using DESeq ^86^. Pathway enrichment analysis along with translation efficiency was performed by iPAGE ^89^. The definition of 5’ TOP mRNAs was adapted as published nanoCAGE data ^90^.

#### Translated intron features

Prediction of intrinsic disorderness was performed using IUPred2A using the short option ^46^. The relative fraction of disordered residues was calculated by counting the total number of disordered amino acid residues (IUPred2A score > 0.5) normalized to the respective length of either translated introns, upstream CDS exons, or CDS. Prediction of low complexicity regions (LCRs) was performed by SEG (http://mendel.imp.ac.at/METHODS/seg.server.html) ^91^.

The hydrophobicity score (Kyte-Doolittle scale) and net charge (Lehninger pKa scale) of the translated introns and their respective upstream exons and CDS were calculated with the Peptides package in R. The relative fraction of amino acids was calculated in R by counting the total number of particular amino acid residues normalized to the respective length of either translated introns, upstream exons or CDS.

Gene ontology enrichment analysis for the genes showing intron translation was performed by Gorilla (Gene Ontology enRIchment anaLysis and visuaLizAtion tool) using the background of HeLa cell expressed genes in our experiment ^92^.

Detailed codes will be available upon request.

### Western blotting

Cells were lysed using the same lysis buffer as used for the ribosome profiling experiment with 1× protease inhibitor cocktail (Roche), omitting cycloheximide. Proteins were transferred to nitrocellulose membranes (Bio-Rad), and the membrane was blocked with Odyssey blocking buffer (TBS) (LI-COR, 927-50000). Anti-4EBP1 [Cell Signaling Technology (CST), 9452], anti-phospho-4EBP1 (Thr37/46) (CST, 2855), anti-phospho-4EBP1 (Ser65) (CST, 9456), anti-S6K1 (49D7) (CST, 2708), anti-phospho-S6K1 (Thr389) (108D2) (CST, 9234), anti-phospho-JNK (Thr183/Tyr185) (CST, 4668), anti-JNK1 (2C6) (CST, 3708), anti-JNK2 (56G8) (CST, 9258), anti-SAPK/JNK (CST, 9252), anti-FLAG (Sigma-Aldrich, F1804), anti-β-actin (MBL, M177-3 and LI-COR, 926-42212), anti-phospho-RAPTOR (S863) (Sigma, SAB1305088), anti-histone H3 (Abcam, ab1791), and anti-SF3B1 (CST, 14434) antibodies were used. IR-dye (680 or 800 nm)-conjugated secondary antibodies (LI-COR, 925-68070/71 and 926-32210/11) were used for detection. Images were collected with the Odyssey CLx Infrared Imaging System (LI-COR). Image studio ver 5.2 (LI-COR) was used for image quantification.

### RT-PCR

HeLa S3 cells were treated with 100 ng/ml SSA, 100 ng/ml PlaB, or MeOH for 6 h. Cells were lysed using ribosome profiling lysis buffer without cycloheximide. Then, total RNA was extracted using TRIzol LS (Thermo Fisher Scientific) and purified by Direct-zol RNA MicroPrep Kits (Zymo Research). Six hundred thirty nanograms of total RNA was annealed to random 9-mer primers (TaKaRa) and reverse-transcribed using ProtoScript II (New England Biolabs). PCR was performed with an equal volume of the acquired cDNA in 25 µl of reaction mixture using PrimeSTAR Max Premix (TaKaRa) and each appropriate primer pair. The primers to detect *DNAJB1* intron retention are listed as follows:

Exon 2-Fw: 5′-GAACCAAAATCACTTTCCCCAAGGAAGG-3′ and Exon 3-Rv: 5′-AATGAGGTCCCCACGTTTCTCGGGTGT-3′.

The PCR conditions were 98°C for 3 min; 35 cycles of 98°C for 10 sec, 52°C for 15 sec, and 72°C for 60 sec; followed by 72°C for 3 min. PCR products were visualized by the MultiNA fragment analyzer (Shimazu) using the DNA-1000 Reagent Kit (Shimazu) and SYBR Gold (Thermo Fisher Scientific).

### BONCAT

HeLa S3 cells (7.5 × 10^6^ cells) were grown 24 h in a 10-cm dish in regular DMEM supplemented with 10% FBS. Medium was removed from the dish and cells were washed by PBS twice. Ten ml of methionine-free DMEM (Invitrogen) was added to the dish and cells were incubated for 30 min at 37°C. The medium was replaced by methionine-free DMEM containing either 100 ng/ml SSA or solvent MeOH along with 50 µM HPG (Jena Bioscience) (3 replicates each). Cells were incubated for 6 h at 37°C. After medium was removed, cells were washed by ice-cold PBS thoroughly and lysed with 500 µl lysis buffer (20 mM Tris-HCl pH 7.5, 150 mM NaCl, 5 mM MgCl_2_, and 1% Triton-X 100). By centrifugation at 20,000 g and 4°C for 10 min, the supernatant and pellet were separated. Pellet fraction was dissolved in lysis buffer along with manual grinding on wall of tube and sonication.

HPG labeled proteins in supernatant and pellet fractions (∼1 mg) were click-conjugated with 50 µM azide-PEG3-biotin (Sigma-Aldrich) by a Click-iT Cell Reaction Buffer Kit (Thermo Fisher Scientific) according to the manufacturer’s instructions. Free azide-PEG3-biotin was removed with MicroSpin G-25 Columns (GE Healthcare). The flow-through fraction was mixed with 300 µl of Dynabeads M280 streptavidin (Invitrogen), which were equilibrated with lysis buffer with 1 mM DTT, and incubated overnight at 4°C with slow rotation. Beads were washed 3 times by lysis buffer with 1 mM DTT). Beads were finally dissolved in bead storage buffer (20 mM Tris-HCl pH 7.5, 150 mM NaCl, and 5 mM MgCl_2_) and LC-MS/MS was performed by on-bead digestion with trypsin (TPCK-treated, Worthington Biochemical).

### LC-MS/MS

After reduction and S-carboxymethylation of the insoluble fraction, the proteins were precipitated by trichloroacetic acid (TCA) (PAGE Clean Up Kit, Nacalai Tesque) and then digested with trypsin (TPCK-treated, Worthington Biochemical). The digestion mixture was separated on a nanoflow LC (Easy nLC 1200, Thermo Fisher Scientific) using a nanoelectrospray ionization spray column (NTCC analytical column; C18, φ75 µm × 100 mm, 3 µm; Nikkyo Technology) with a linear gradient of 0-40% Buffer B (80% acetonitrile and 0.1% formic acid) in Buffer A (0.1% formic acid) and a flow rate of 300 nl/min over 220 min, coupled online to a Q-Exactive HFX mass spectrometer (ThermoFisher Scientific) that was equipped with a nanospray ion source. The mass spectrometer was operated in positive-ion mode, and MS and MS/MS spectra were acquired with a data-dependent TOP 10 method. Proteins were identified and quantified using Proteome Discoverer 2.2 (Thermo Fisher Scientific) with MASCOT program ver 2.6 (Matrix science) using an in-house database.

### DNA constructs

#### pCDNA5/FRTTO-FTH1, FTH1*, RAB32, RAB32*, FADD, FADD*, IRF2BP2, IRF2BP2*, p27, and p27*

DNA fragments containing the first coding exon and following the intron until the first in-frame stop codon from *FTH1*, *RAB32*, *FADD*, *IRF2BP2*, or *p27* (*CDKN1B)* were PCR-amplified from the HEK cell genome. DNA fragments coding the *FTH1*, *RAB32*, *FADD*, *IRF2BP2*, or *p27* (*CDKN1B)* CDS were PCR-amplified from HEK cell cDNA. An N-terminal 1× FLAG tag was inserted in frame upstream during PCR amplification. These PCR products were inserted into a pcDNA5/FRT/TO vector (Invitrogen) via the HindIII and BamHI sites by Gibson assembly (New England Biolabs). The sequences of the final constructs were verified by plasmid sequencing.

#### pColdI-mCherry-FTH1 and FTH1*

DNA fragments for FTH1 and FTH1* were PCR-amplified from the HEK cell cDNA and genome, respectively, and inserted into a pColdI vector (TaKaRa) with mCherry-tag by Gibson Assembly (New England Biolabs).

#### psiCHECK2-ATP5O

The non-TOP 5′ UTR sequence of ATP5O (5′-CGGGAGAAG-3′) was identified in published nanoCAGE data ^90^ and inserted between the T7 promoter and *Renilla* luciferase in psiCHECK2 (Promega) by Gibson Assembly.

#### psiCHECK2-HCV-FL

*The* DNA region spanning from *Renilla* luciferase to the HSV-TK promoter was removed from psiCHECK2-HCV IRES ^67^ by PCR and Gibson Assembly.

### DNA transfection

Transfection of 1 µg of each individual DNA was performed in six-well plates using FuGENE HD (Promega) for HeLa S3 cells according to the manufacturer’s instructions.

### Recombinant protein purification and imaging

*E. coli* BL21 Star (DE3) cells (Thermo Fisher Scientific) were transformed with pColdI-mCherry-FTH1 or FTH1* and cultivated to OD_600_ ∼0.5 in 1 l of LB with ampicillin. The protein was inducted with 1 mM IPTG at 15°C overnight. The collected cell pellet was flash-frozen by liquid nitrogen and stored at -80°C.

The pellet was resuspended in Buffer A (20 mM HEPES-NaOH pH 7.5, 1 M NaCl, 10 mM imidazole, 0.5% NP-40, and 10 mM 2-mercaptoethanol) and then sonicated on ice. The cell lysate was centrifuged at 10,000 g for 20 min at 4°C. The supernatant was incubated with 1.5 ml bed volume of Ni-NTA Agarose (Qiagen) equilibrated with Buffer C. Then, beads were washed with Buffer D (20 mM HEPES-NaOH pH 7.5, 1 M NaCl, 20 mM imidazole, and 10 mM 2-mercaptoethanol) on a gravity column (Bio-Rad). The bound proteins were eluted with elution buffer (20 mM HEPES-NaOH pH 7.5, 1 M NaCl, 250 mM imidazole, 10 mM 2-mercaptoethanol, and 10% glycerol). Using an NGC chromatography system (Bio-Rad), the proteins were loaded into HiLoad 16/60 Superdex 75 pg (GE Healthcare) in Buffer E (20 mM HEPES-NaOH pH 7.5, 1 M NaCl, and 1 mM DTT) and purified according to the size of the proteins. The proteins were concentrated using Vivaspin 6, 10,000 MWCO (Sartorius), flash-frozen by liquid nitrogen, and stored at -80°C.

Microscopic images of the recombinant proteins in chambered cover glass (Thermo Fisher Scientific, Nunc Lab-Tek II) were taken using a confocal microscope (Olympus, FV3000). The proteins were diluted in 20 mM HEPES-NaOH pH 7.5, 200 mM NaCl, and 1 mM DTT.

### Immunostaining and microscopic analysis

Cells were grown on coverslips, transfected with plasmids as described above, washed with PBS, fixed with 4% paraformaldehyde for 45 min, permeabilized with 0.2% Triton-X 100 in PBS, incubated with 10% normal goat serum (NGS) (Thermo Fisher Scientific, PCN5000) diluted in 0.2% Triton-X 100 in PBS as the blocking solution for 45 min, and then incubated with the anti-FLAG antibody (Sigma-Aldrich, F1804) in the blocking solution at 4°C overnight.

The next day, cells were washed three times with 0.2% Triton-X 100 in PBS followed by incubation with the secondary antibody anti-Alexa 594 (Thermo Fisher Scientific, A11005) or anti-Alexa 488 (Thermo Fisher Scientific, A11008) in blocking solution for 50 min. Washing was performed three times for 10 min each with 0.2% Triton-X 100 including Hoechst nuclei stain (Hoechst 33342, Invitrogen, H3570) during the second wash. Coverslips were mounted on microscope slides using ProLong Diamond Antifade Mountant (Thermo Fisher Scientific, P36965). Fluorescence imaging was obtained with a DeltaVision imaging system (Applied Precision) and deconvolved using SoftWoRx 5.5 (Applied Precision). Automated quantification of GFP fluorescence intensity and size of GFP aggregates were obtained using ImageJ. For the experiment with SSA treatment, immunostaining was not performed except for Hoechst staining.

### Proteotoxic stress reporter transfection and luciferase activity assay

HeLa S3 cells (∼40,000) were seeded in 24-well plates in 1 ml DMEM in triplicate. The next day, transfection of 0.5 µg of either pCI-neo Fluc-EGFP (Addgene plasmid #90170; http://n2t.net/addgene:90170; RRID:Addgene_90170) or pCI-neo FlucDM-EGFP (Addgene plasmid #90172; http://n2t.net/addgene:90172; RRID:Addgene_90172), both gifts from Franz-Ulrich Hartl ^60^, was carried out using FuGENE HD (Promega) according to the manufacturer’s instructions.

For the firefly luciferase activity assay, cells were washed with PBS and lysed with 1× passive lysis buffer (Promega). Luciferase assay reagent and GLOMAX (both Promega) were used to detect luminescence according to the manufacturer’s instructions. The relative luciferase activity was determined by normalizing the firefly luciferase activity by the respective total EGFP protein amount quantified by Western blot using a GFP antibody (Clontech, 632460) in Image Studio ver 5.2 (LI-COR).

### *In vitro* mTOR kinase assay

mTOR kinase activity was monitored by LANCE Ultra time-resolved fluorescence resonance energy transfer (TR-FRET, PerkinElmer) following the manufacturer’s instructions. mTOR enzyme (10 nM), ATP (90 µM), and ULight-S6K1 (Thr389) peptide (25 nM) were incubated in kinase buffer (50 mM HEPES pH 7.5, 1 mM EGTA, 3 mM MnCl_2_, 10 mM MgCl_2_, 2 mM DTT, and 0.01% Tween 20) at room temperature for 1 h in white 384-well opti-plates. The reaction was stopped by adding EDTA to a final concentration of 10 mM. A europium-labeled anti-phospho-S6K1 (Thr389) antibody was then added to the reaction at 2 nM to detect the phosphorylated peptide. The signal intensity of the emitted light was measured with an EnVision Multilabel reader (PerkinElmer) in TR-FRET mode (excitation at 320 nm and emission at 665 nm).

### RNA interference

ON-TARGETplus siRNAs against Nontargeting #1 (D-001810-01-05), human SF3B1 siRNA (L-020061-01-0005), human JNK1 (L-003514-00-0005), and human JNK2 (L-003505-00-0005) were obtained from Dharmacon. For the knockdown experiments, 2 × 10^5^ HeLa S3 cells per well were seeded in 6-well plates. The next day, 30 pmol of siRNA was transfected into cells using Lipofectamine RNAiMAX Transfection Reagent (Thermo Fisher Scientific) according to the manufacturer’s manual. Thirty-six hours after transfecting the SF3B1 siRNAs, cells were lysed as mentioned above in the Western blotting section. In the case of JNK knockdown experiments, 42 h after transfection, cells were treated with SSA for 6 h and then lysed.

### Nascent peptide labeling by OP-puro

Nascent peptides were labeled with 20 µM OP-puro (Jena Bioscience) in 24-well plates for 30 min after 5.5 h of SSA or MeOH challenge. Cells were washed with ice-cold PBS and lysed with 60 µl buffer (20 mM Tris-HCl pH 7.5, 150 mM NaCl, 5 mM MgCl_2_, and 1% Triton-X 100) and centrifuged at 20,000 g and 4°C for 10 min. Supernatants were used for nascent peptide labeling with 50 µM azide-conjugated IR-800 dye (LI-COR) by a Click-iT Cell Reaction Buffer Kit (Thermo Fisher Scientific) according to the manufacturer’s instructions and were run on SDS-PAGE. Images were acquired on an Odyssey CLx Infrared Imaging System (LI-COR) for the detection of nascent peptides at an IR 800 nm signal. The total proteins were measured by staining with Coomassie Brilliant Blue (CBB) (Wako chemicals), giving a signal at IR 700 nm. The images were quantified with Image Studio ver 5.2 (LI-COR).

### Reporter mRNA preparation

For *Renilla* luciferase reporter mRNAs, DNA fragments for *in vitro* transcription were PCR-amplified from psiCHECK2-EIF2S3 ^67^ and psiCHECK2-ATP5O as templates using the following primers: 5′-TAATACGACTCACTATAGG-3′ and 5′-CTGTGTGTTGGTTTTTTGTGTGTG-3′. *In vitro* transcription, capping and polyadenylation of the reporter RNA were performed with the T7-Scribe Standard RNA IVT Kit, ScriptCap m^7^G Capping System, ScriptCap 2’-O-methyltransferase, and A-Plus Poly(A) Polymerase Tailing Kit (CELLSCRIPT) according to the manufacturer’s protocol.

For the HCV-firefly luciferase mRNA reporter, the DNA fragment was PCR-amplified from psiCHECK2-HCV-FL with the following primers: 5′-TGACTAATACGACTCACTATAGG-3′ and 5′-TGTATCTTATCATGTCTGCTCGAAG-3′.

AP_3_G (A-cap) (Jena Bioscience) was added to the *in vitro* transcription reaction, skipping the capping reaction.

### Luciferase reporter assay

HeLa S3 cells (1 × 10^5^) were seeded onto 24-well plates in triplicate. Two hours after 100 ng/ml SSA treatment, 0.02 µg of each mRNA reporter per well was transfected the next day using the TransIT-mRNA Transfection Kit (Mirus) according to the manufacturer’s instructions. Four hours after transfection, cells were washed with PBS and lysed with 1× passive lysis buffer (Promega). The Dual-Luciferase Reporter Assay System and GLOMAX (both Promega) were used to detect luminescence according to the manufacturer’s instructions.

### 7-Methyl-guanosine (m^7^G) pulldown assays

HeLa S3 cells were grown until 70-80% confluency in a 15-cm dish. Following 10 h of 100 ng/ml SSA treatment, HeLa S3 cells were washed with ice-cold PBS and lysed with 1200 µl hypotonic lysis buffer [10 mM HEPES-NaOH pH 7.5, 10 mM KCl, 1.5 mM MgCl_2_, 1 mM DTT, and 1× protease inhibitor cocktail (Nacalai)]. After centrifugation at 20,000 g and 4°C for 10 min, the lysate was precleared with 150 µl of Pierce control agarose resin (Thermo Fisher Scientific) equilibrated with hypotonic lysis buffer at 4°C for 1 h. The precleared lysate was incubated with 40 µl of agarose (blank) (Jena Bioscience) or γ-aminophenyl-m^7^GTP (C10-spacer)-agarose (Jena Bioscience) equilibrated with hypotonic wash buffer [10 mM HEPES-NaOH pH 7.5, 10 mM KCl, 1.5 mM MgCl_2_, 1 mM DTT, 0.02% Triton-X 100, and 50 µg/ml tRNA from baker’s yeast (Sigma-Aldrich)] at 4°C for 1 h and then washed with hypotonic wash buffer 3 times. Proteins were eluted with LDS sample buffer (Thermo Fisher Scientific) at 100°C for 4 min and examined by Western blotting.

## Accession numbers

The ribosome profiling and RNA-Seq data (GSE129305) used in this study were deposited in NCBI.

---

**Extended Data Fig. 1:**
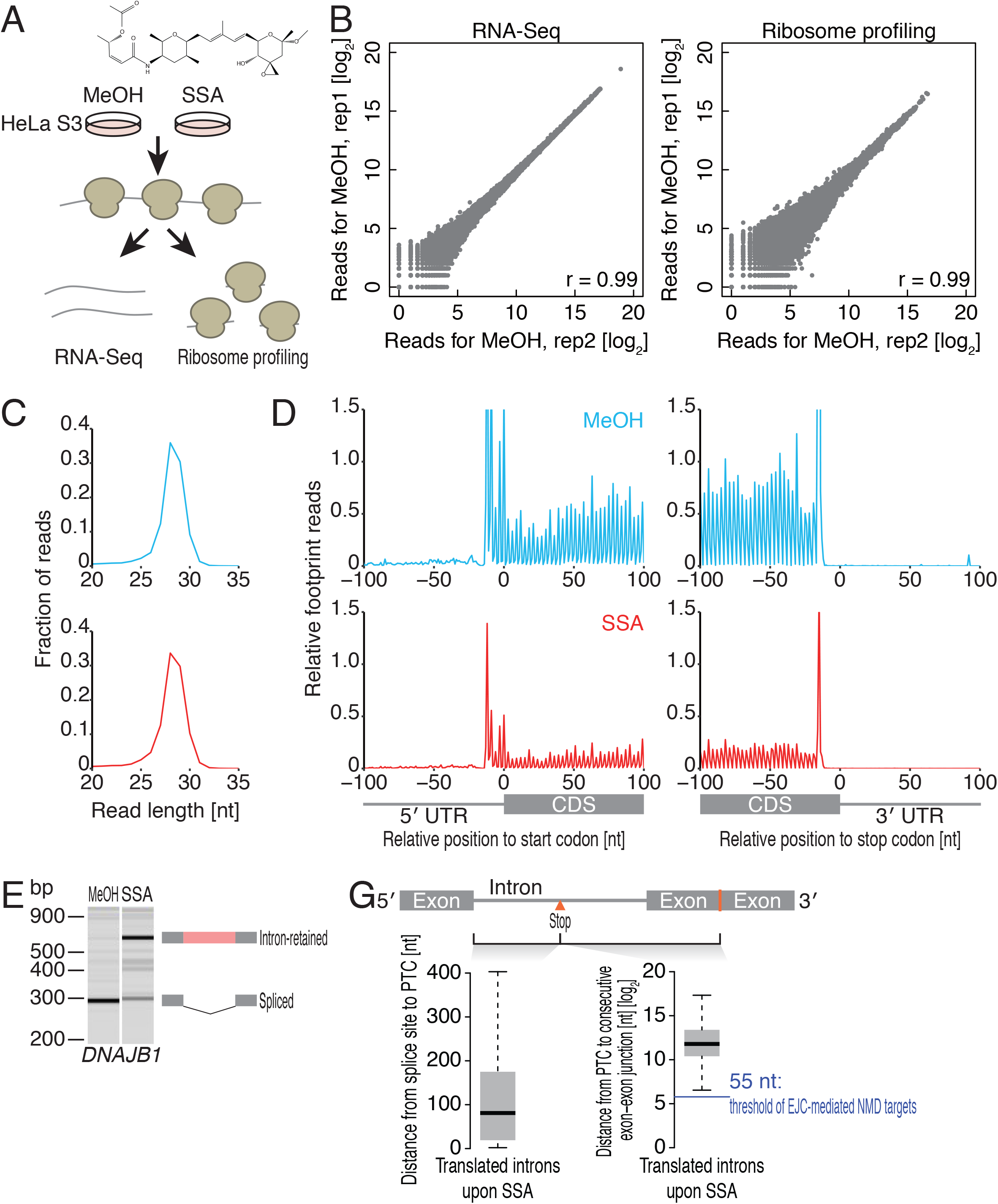
Genome-wide analysis of RNA and translation upon splicing inhibition by SSA. (A) Schematic presentation of the experimental design. HeLa S3 cells were subjected to solvent (MeOH) or SSA (100 ng/ml) treatment for 6 h before lysis. RNA-Seq and ribosome profiling libraries were prepared from the same samples. (B) The RNA-Seq (left panel) and ribosome profiling (right panel) experiments were shown to have high reproducibility by a high Pearson’s correlation coefficient (r) between two independent experiments under vehicle treatment. (C) The size distribution of ribosome footprint reads. (D) Meta-gene analysis of ribosomal footprint density for MeOH and SSA around the start codon (left panel) and stop codon (right panel). The 5′ ends of the reads are depicted. Reads are normalized to the sum of mitochondrial footprint reads. (E) RT-PCR analysis to detect spliced and unspliced forms of *DNAJB1* upon splicing inhibition by SSA using primers spanning exons 2 and 3. (G) Boxplot showing the distribution of length from the 5’ splice site to the PTC (left) and the PTC to the next exon-exon junction (right). The threshold for EJC-mediated NMD targets (55 nt) is drawn in blue.

**Extended Data Fig. 2:**
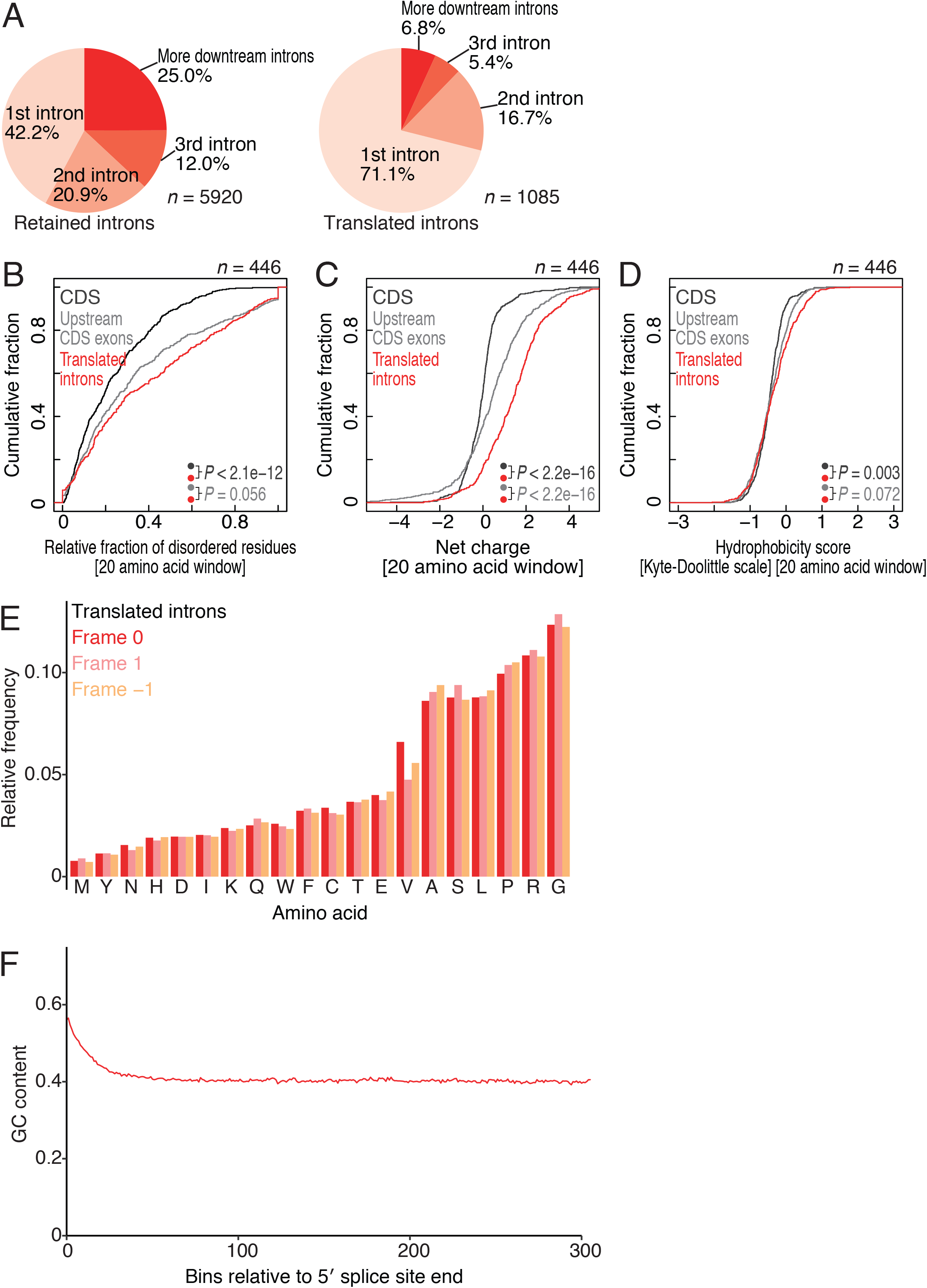
Characterization of SSA-mediated translated introns. (A) Fraction of annotated intron position in which RNA-seq (left) and ribosome profiling (right) reads were obtained. (B-D) Cumulative distribution of relative disorderness (IUPred2A score) (B), net charge (Lehninger pKa scale) (C), and hydrophobicity score (Kyte-Doolittle scale) (D) of amino acid residues in translated introns, upstream CDS exons, and full CDS. Twenty amino acid windows were considered. The significance was calculated by Wilcoxon’s test. (E) Relative frequency of amino acids present in translated introns. The predicted relative amino acid frequencies of the other two frames (frames -1 and +1) were included. (F) A plot showing the average GC content for retained introns in every 30 nt window from the 5′ splice site towards the 3′ splice site.

**Extended Data Fig. 3:**
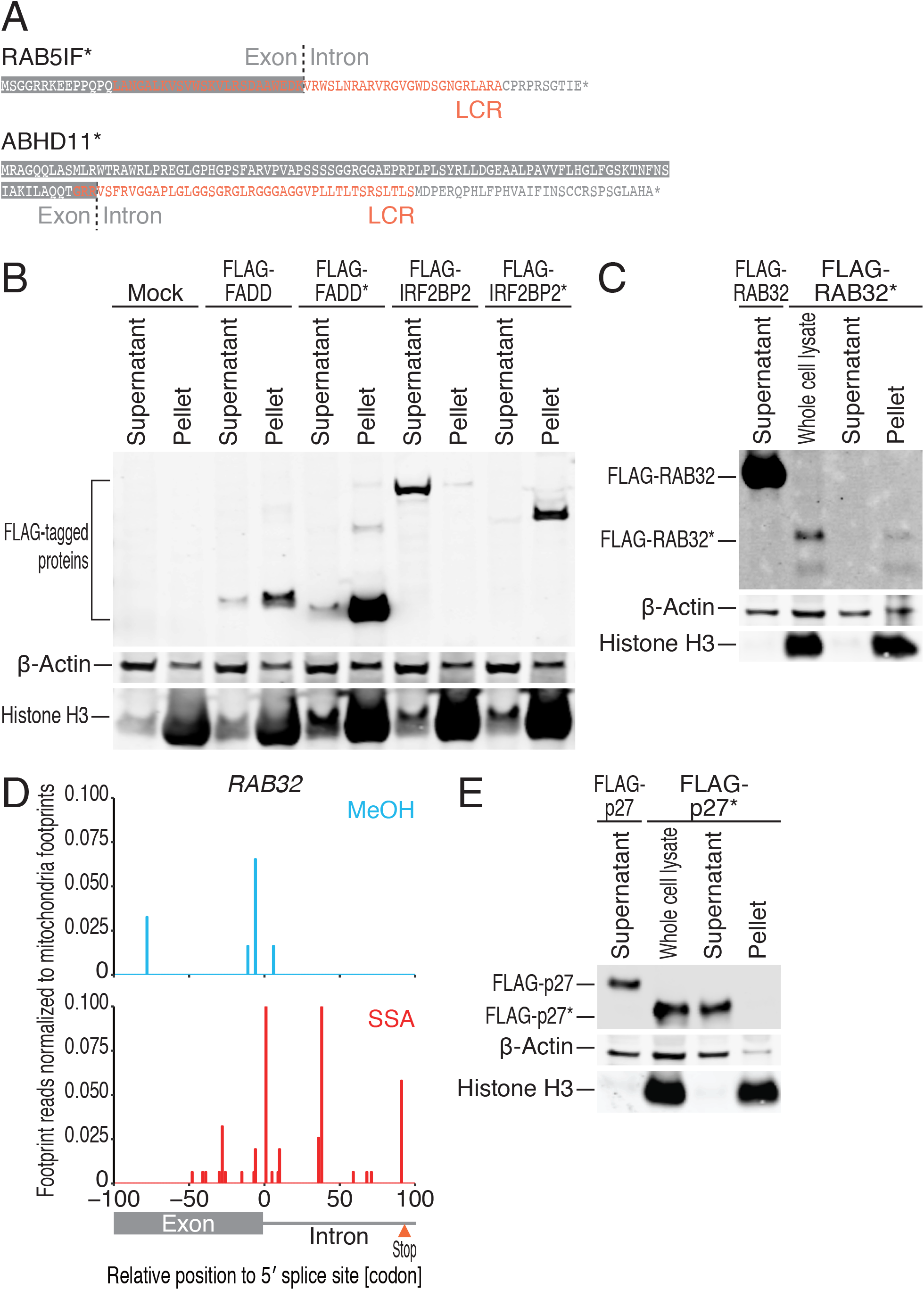
Characterization of intron-derived peptides. (A) LCR found in the condensate-prone translated introns (Fig. 3A). (B and C) Western blot of FLAG-tagged proteins ectopically expressed in HeLa S3 cells. Cell lysate was subfractionated by centrifugation. (D) Ribosome footprint accumulation in introns of *RAB32* under splicing inhibition. Reads were normalized to the sum of mitochondrial footprint reads. (E) Western blot of ectopically expressed FLAG-tagged p27* in HeLa S3 cells. Cell lysate was subfractionated by centrifugation.

**Extended Data Fig. 4:**
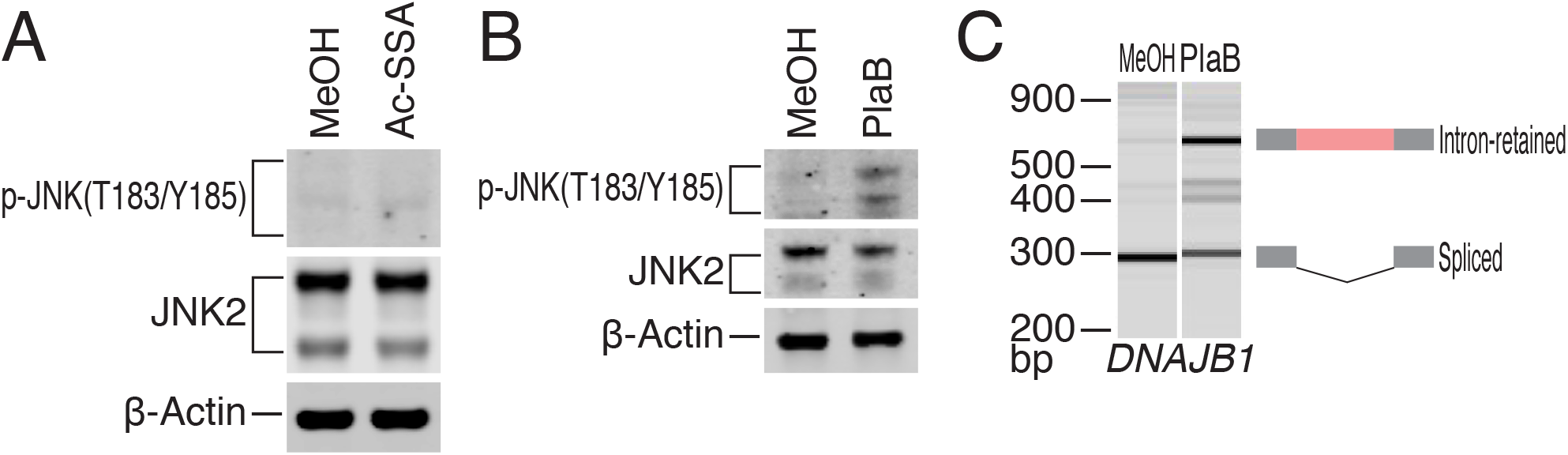
Characterization of JNK activation upon compounds. (A and B) HeLa S3 cells were either treated with 100 ng/ml Ac-SSA (A) or 1 µg/ml PlaB (B) for 10 h, and the cell lysates were immunoblotted with the indicated antibodies. (C) RT-PCR analysis to detect spliced and unspliced forms of *DNAJB1* using primers spanning exons 2 and 3 upon splicing inhibition by PlaB.

**Extended Data Fig. 5:**
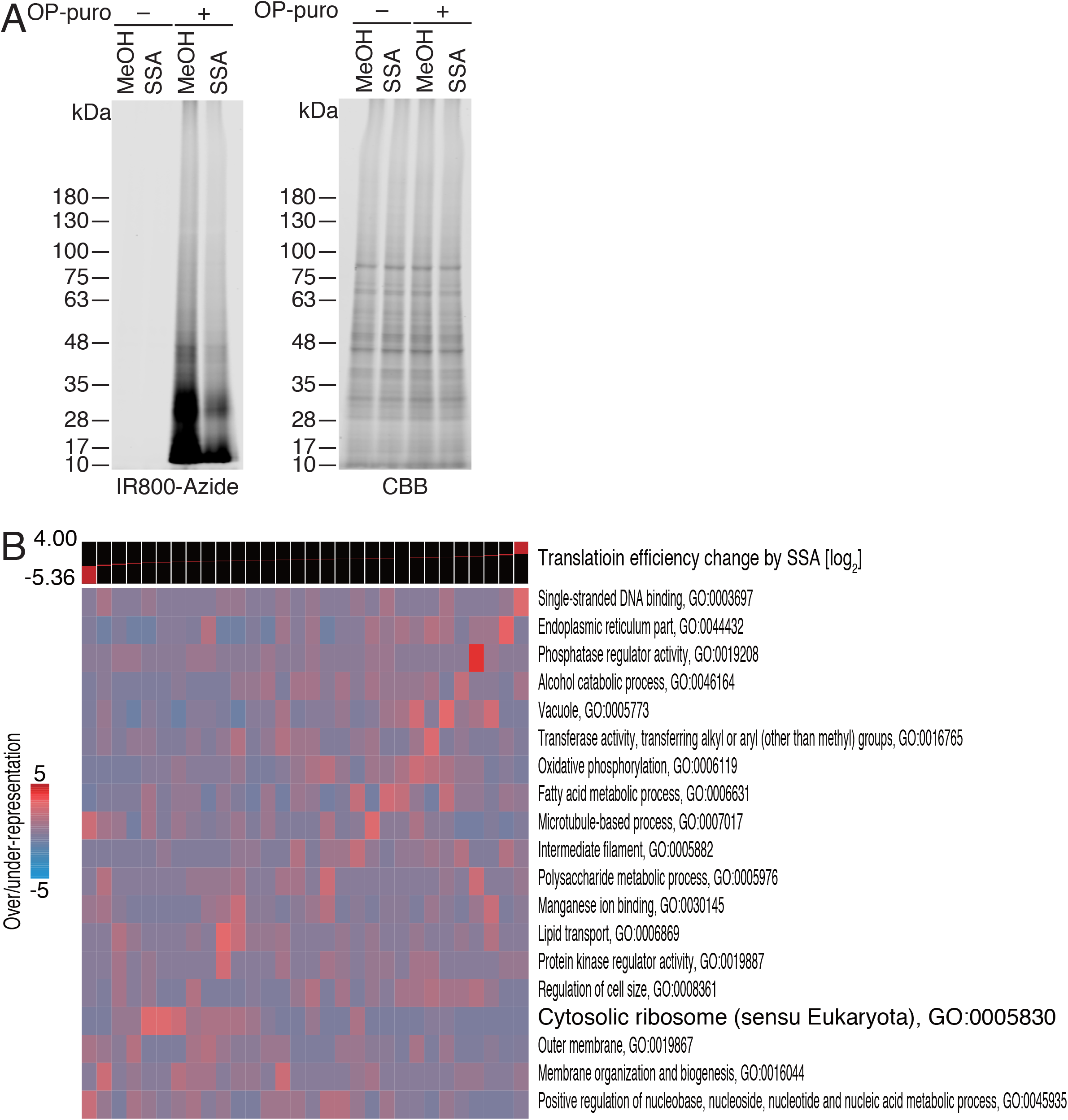
Characterization of translation change induced by SSA treatment. (A) Change in translation activity upon SSA treatment in HeLa S3 cells monitored by nascent peptide labeling with OP-puro. Nascent peptides labeled with OP-puro were labeled with IR-800 dye by click reaction and run on SDS-PAGE (left). The total protein amount was quantified by CBB staining. (B) GO pathway analysis based on translation efficiency change during SSA treatment, visualized with iPAGE ^89^.

**Extended Data Fig. 6:**
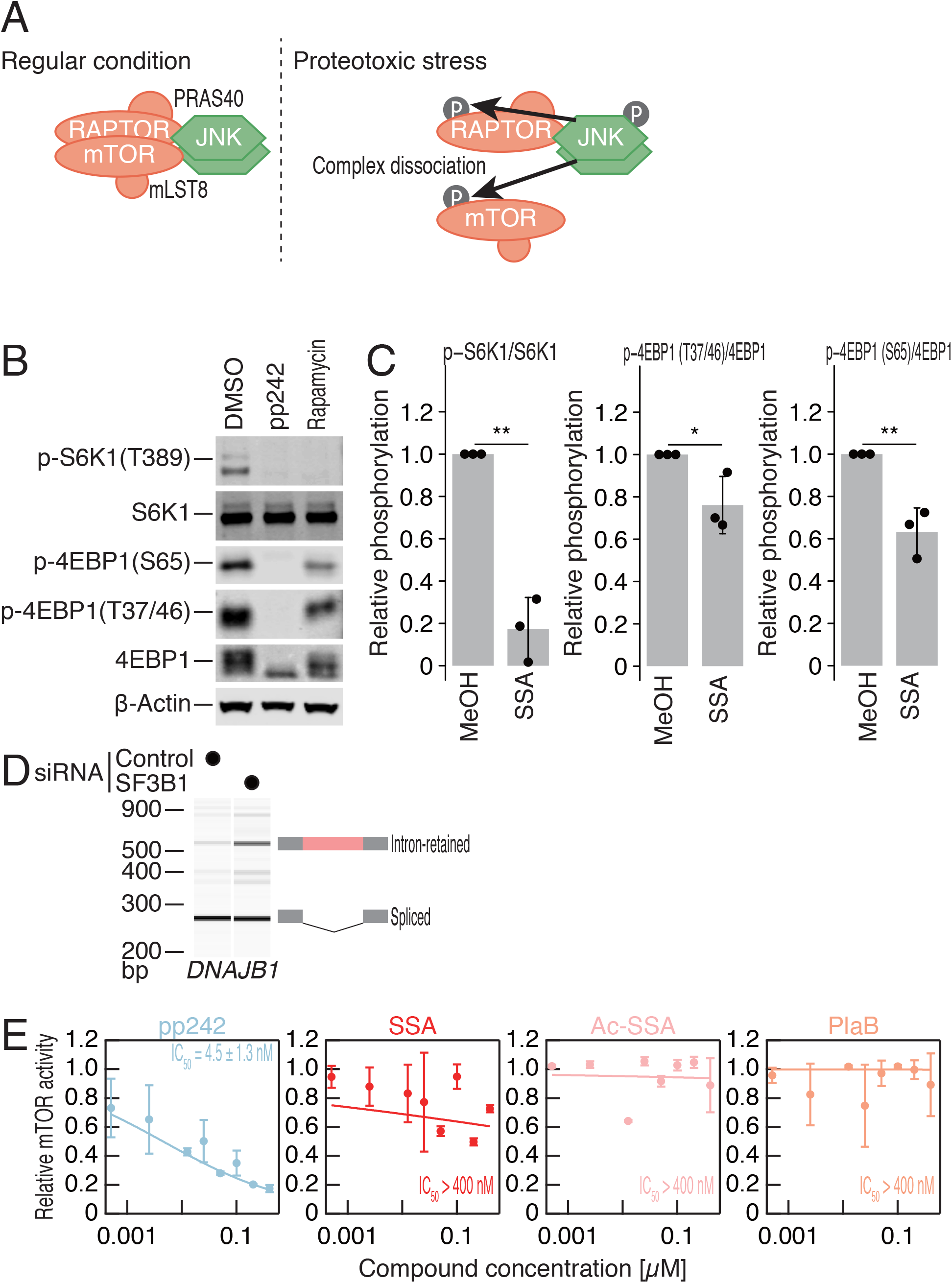
Characterization of SSA-mediated mTORC1 inhibition. (A) Schematic of proteotoxic stress-mediated mTORC1 inhibition via activated JNK. (B) HeLa S3 cells were treated with DMSO solvent, 2.5 µM pp242 (ATP-competitive mTOR inhibitor), or 100 ng/ml rapamycin for 10 h. Representative Western blots for either phosphorylated or bulk S6K1 and 4EBP1 are shown. (C) Western blots for phosphorylated and total S6K1 and 4EBP1 were quantified upon 10 h treatment of solvent MeOH and SSA . The data represent the mean and s.d. (n = 3). *P* values were calculated using Student’s *t*-test (one-tailed). *, *P* < 0.05; **, *P* < 0.01. (D) RT-PCR analysis to detect spliced and unspliced forms of *DNAJB1* using primers spanning exons 2 and 3 upon splicing inhibition by knockdown of SF3B1. (E) *In vitro* mTOR kinase assay with pp242, SSA, Ac-SSA, and PlaB. IC_50_ values were calculated for each of the compounds.

**Extended Data Fig. 7:**
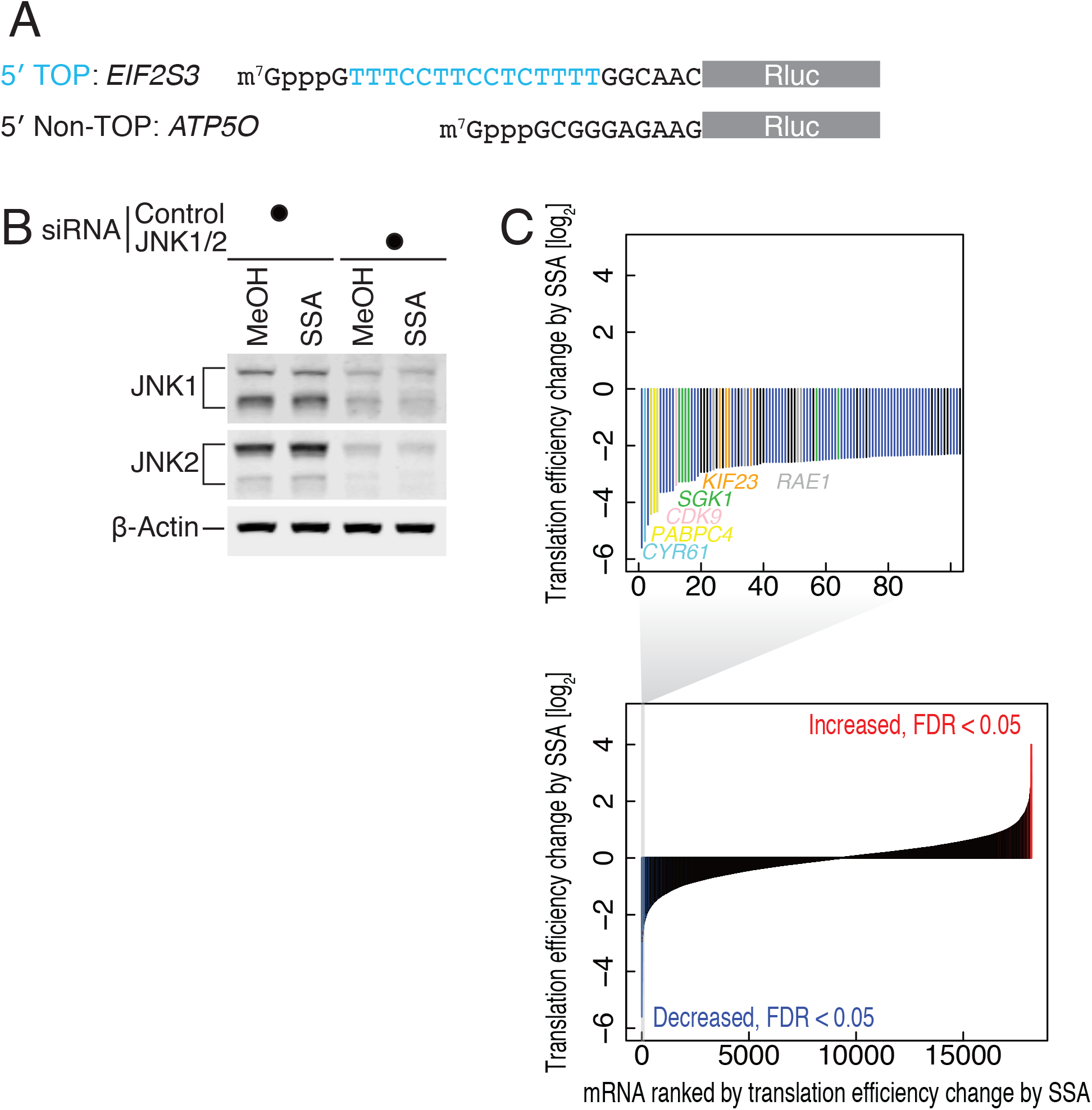
Translation efficiency changes of tumor-related genes in ribosome profiling. (A) The sequence of 5′ UTRs used in the reporters for Fig. 7C and D. (B) JNK1 and JNK2 were simultaneously knocked down in HeLa S3 cells before treatment with 100 ng/ml SSA for 10 h. The cell lysates were immunoblotted with the indicated antibodies. (C) Rank plot of translationally downregulated genes. Cancer-related genes are highlighted.

**Extended Data Table 1: List of chimeric intron protein sequences and characteristics.** Each intron is listed with its UCSC identifier and individual scores (IUPred2A-based disordered fraction, hydrophobicity and net charge). Either a 10 or 20 amino acid window was considered.

**Extended Data Table 2: List of chimeric intron proteins depicted by LC-MS/MS.**

Each intron is listed with its UCSC identifier. The respective names of mRNA and gene for the intron are listed.

